# A new level of RNA-based plant protection - dsRNAs designed from functionally characterized siRNAs highly effective against Cucumber Mosaic Virus

**DOI:** 10.1101/2024.06.03.597145

**Authors:** Marie Knoblich, Torsten Gursinsky, Selma Gago-Zachert, Claus Weinholdt, Jan Grau, Sven-Erik Behrens

**Affiliations:** Institute of Biochemistry and Biotechnology, Martin Luther University Halle-Wittenberg, Charles Tanford Protein Centre, Kurt-Mothes-Str. 3A, 06120 Halle (Saale); Institute of Computer Science, Martin Luther University Halle-Wittenberg, Von-Seckendorff-Platz 1, 06120 Halle (Saale)

**Keywords:** dsRNA, RNA silencing, RNAi, crop protection, siRNA, ASO

## Abstract

RNA-mediated crop protection increasingly becomes a viable alternative to agrochemicals that threaten biodiversity and human health. Pathogen-derived double-stranded dsRNAs are processed into small interfering RNAs (siRNAs), which can then induce silencing of target RNAs, *e.g.* viral genomes. However, with currently used dsRNAs, which largely consist of undefined regions of the target RNAs, silencing is often ineffective: processing generates siRNA pools that contain only a few functionally effective siRNAs (here called *e*siRNAs). Using a recently developed *in vitro* screen that reliably identifies *e*siRNAs from siRNA pools, we identified *e*siRNAs against Cucumber Mosaic Virus (CMV), a devastating plant pathogen. Topical application of *e*siRNAs to plants resulted in highly effective protection against massive CMV infection. However, optimal protection was achieved with newly designed multivalent “effective dsRNAs” (*e*dsRNAs), which contain the sequences of several *e*siRNAs and are preferentially processed into precisely these *e*siRNAs. The *e*siRNA components can attack one or more target RNAs at different sites, be active in different silencing complexes and provide cross-protection against different viral variants, important properties for combating rapidly mutating pathogens such as CMV. *e*siRNAs and *e*dsRNAs have thus been established as a new class of “RNA actives” that significantly increase the efficacy and specificity of RNA-mediated plant protection.

## Introduction

Virus-induced plant diseases remain a major problem in agriculture, recently exacerbated by global trade and climate change (1,2). The most common method of controlling viral infections is the extensive use of chemical pesticides, which target the vectors but often have a non-specific effect on arthropods and can be harmful to humans (3). Urgently needed alternative crop protection methods should not only be environmentally sustainable, but also specific, *i.e.* effective only against a specific target pathogen, and adaptable to the evolution of the pathogen. One strategy to meet these complex demands is to trigger the RNA silencing component of the plant’s immune response against the pathogen (4,5).

RNA silencing is a conserved cellular defense mechanism, best characterized in higher eukaryotes, that serves to block (silence) or modulate gene expression at the level of target RNAs. Antiviral RNA silencing is triggered by double-stranded (ds) regions of RNA molecules, particularly by nearly completely double-stranded replication intermediates of positive-strand RNA viruses, the most common class of viral pathogens in plants. Intracellular dsRNA is perceived by Dicer-like proteins (DCLs), which belong to the type III endonuclease family (6). Of the four known DCLs in the model plant *Arabidopsis thaliana*, DCL4 and DCL2 are central to antiviral RNA silencing (7,8). In a process whose molecular details are not yet fully understood, DCL4 and DCL2 hydrolyze dsRNAs into 21 nucleotide (nt)-long or 22 nt-long small interfering RNAs (siRNAs), *i.e.,* RNA duplexes that are phosphorylated at their 5’-end and have a 2 nt single-stranded overhang with 3’-hydroxyl groups at their 3’-end (6,9). The siRNAs become active in RNA-induced silencing complexes (RISCs). The major components of RISCs are Argonaute (AGO) endonucleases, of which AGO1 and AGO2 have been shown to be significantly involved in antiviral RNA silencing (10). After binding of the siRNA duplex to AGO, one strand, the guide strand, remains bound while the other, the passenger strand, is degraded (11); AGO1 and AGO2 preferentially incorporate guide strands containing a 5’-terminal U or A nucleotide, respectively (12,13). The complementarity of the bound guide strands to the target RNAs is central to the activity of RISC; it is naturally highest for siRNAs that were originally processed by the DCLs from these target RNAs. Association of the RISC with the target RNA then enables AGO-catalyzed endonucleolytic hydrolysis (“slicing”) between the nucleotides of the target RNA that are opposite nucleotides 10 and 11 of the siRNA guide strand strand (14). RNA silencing directed by siRNAs can thus result in temporary inhibition of gene expression (4), and RISC-mediated slicing of viral RNAs can further enhance the silencing response by inducing the biogenesis of secondary siRNAs *via* a process that involves host-encoded RNA-dependent RNA polymerases (RDRs) and DCLs (7,15-17) (**Supplementary Figure 1A**).

Virus-derived dsRNAs or siRNAs are already used as antivirals in crop protection. Several plant varieties have been developed that transgenically produce dsRNA and are thus protected against infection with the corresponding viral pathogen (18,19). However, the production of transgenes is time-consuming and costly, and their release is prohibited in many countries due to safety concerns. Furthermore, the plants are potentially susceptible to infection by variants of the pathogen against which they were bred for resistance (20). Alternatively, there are encouraging reports suggesting that dsRNA suspensions, in the simplest case in the form of a spray (21), can be used for topical/transient antiviral applications in plants (22). The prospects for success of these approaches are strengthened by significant advances in the production of dsRNA on a gram or even kilogram scale (23), as well as reports suggesting that certain formulations may stabilize the biodegradable RNAs and facilitate their uptake by the plant (24–28).

However, using “RNA actives”, as we refer to them here, in topical applications is still at an early stage of development for two main reasons. First, the transfer of RNA into plant tissues, particularly the phloem, where the life cycles of several viruses are restricted, remains a major technical challenge (29). The second reason specifically addressed in this study is that both natural and artificially induced antiviral RNA-silencing processes are inefficient. Several independent observations suggest that this is mainly due to the fact that although DCLs generate a large number (a “pool”) of siRNAs from target RNAs, very few of them support AGO-catalyzed cleavage (30–32). One hypothesis that could explain this is that target RNAs are highly structured and that the association of guide strand/AGO/RISC by complementary base pairing, and thus effective silencing, is only possible in a few regions of the RNA (33,34), which are referred to here as “accessible sites”, or a-sites (35). This is particularly relevant when considering the structures of viral genomic RNAs, which have evolved to enable translation and replication, but also to evade RNA silencing by the plant host. Virus-derived siRNAs can even act as decoys by saturating AGO/RISC or silencing cellular targets, facilitating infection (31,36–39).

To date, artificial dsRNAs consisting of large portions of viral genomes or mRNAs have been used in both transgenic and topical RNA-silencing approaches. It is clear from the above that such dsRNAs have the same application problems as viral RNAs themselves: DCLs generate a large pool of siRNAs, only a few of which are actually active. In addition, there is a risk that pool-derived siRNAs may cause “off-target effects”, *i.e.*, the silencing of non-target RNAs, *e.g.,* by base-pairing of guide strands that are not fully complementary but still sufficient to mediate cleavage (40,41).

A globally distributed plant virus of major economic importance is Cucumber mosaic virus (CMV; family *Bromoviridae*). CMV is transmitted by about 90 aphid species and infects more than 1200 plant species, including a large number of agricultural crops. Symptoms of CMV infections include systemic mosaic symptoms, leaf deformation, systemic necrosis, chlorosis, dwarfism, and fruit damage. Apparently, the virus can persist in the seed over winter and cause primary infections early in the growing season (42–45). Several CMV-resistant plant species, including tobacco, cucumber, tomato, melon, squash and pepper, have been generated by transgenic approaches (46) using viral dsRNA-expressing constructs, and promising protective effects have also been observed with topical applications of dsRNAs (47–50). The CMV genome consists of three positive-strand RNA segments (RNA 1, RNA 2, RNA 3; **Supplementary Figure 1B**) that are packaged individually into capsids and cause successful infection when transmitted together. As a result, CMV is capable of reassortment resulting in high mutation rates and antigenic shifts. Transgenic plants are therefore at high risk of loss of resistance, and topical applications require particularly high levels of efficacy.

In previous studies, we and others have established cytoplasmic extracts of cultured *Nicotiana tabacum* BY-2 cells, called BY-2 lysates (BYL), as a versatile experimental tool (51,52). BYL supports the *in vitro* translation of (exogenously added) mRNAs into proteins. Most importantly, BYL recapitulates the key steps of primary RNA silencing: *i.e.*, exogenously added dsRNAs are processed by BYL-endogenous DCLs (32,53), and functional RISC can be reconstituted with an *in vitro-*translated AGO protein and siRNAs (54,55). The activity of the *in vitro*-generated RISC can be tested in a “slicer assay” that detects AGO-mediated cleavage of a target RNA (54–57) (**Supplementary Figure 1C**). Using the BYL system, we have developed a simple *in vitro* experimental method called the “*e*NA screen” to identify nucleic acids, such as siRNA guide strands or DNA antisense oligonucleotides (ASO), which can bind to the a-sites of target RNAs and enable efficient endonuclease-catalyzed hydrolysis (32,35). From a DCL-generated siRNA pool, *e*NA screening reliably identifies those siRNAs, referred to here as “*e*siRNAs”, that cause efficient AGO/RISC-mediated cleavage of a selected target RNA.

By applying the *e*NA screen to RNAs 2 and 3 of CMV strain Fny, we have identified *e*siRNA*s* that are highly protective against infection by the virus in topical plant protection experiments. Based on these functionally characterized *e*siRNAs, we have developed “*e*dsRNA” constructs that, when processed by DCLs, produce high levels of these same *e*siRNAs and thus provide significantly better protection against CMV infection than a comparable conventionally organized dsRNA. The tools established here will help to overcome the lack of efficacy of natural RNA silencing and to provide rapidly and reliably *e*siRNAs and *e*dsRNAs that could be flexibly used in future topical applications against CMV and other plant pathogens.

## Materials and Methods

### Cell culture and preparation of BYL

*Nicotiana tabacum* BY-2 cells (58) were cultivated at 23 °C in Murashige & Skoog liquid medium (Duchefa Biochemie, Haarlem, The Netherlands, #M0221.0025). Protoplasts were generated by treatment with Cellulase RS and Pectolyase Y-23 (Duchefa Biochemie, #C8003 and #P8004) and cytoplasmic extract (BY-2 lysate, BYL) was prepared after evacuolation of protoplasts by percoll gradient centrifugation (Cytiva, Uppsala, Sweden, #17089102) as previously described (51,52).

### Oligonucleotides and siRNAs

DNA oligonucleotides were purchased from Eurofins Genomics (Ebersberg, Germany). The siRNA gf698 targeting GFP mRNA was described earlier (54,55). Single-stranded RNA oligonucleotides were synthesized by Biomers (Ulm, Germany). Sequences of DNA and RNA oligonucleotides are listed in **Supplementary Table S1**. To produce siRNA duplexes, both strands were mixed in siRNA annealing buffer (30 mM Hepes/KOH, pH 7.4, 100 mM KOAc, 2 mM MgOAc) and incubated for 1 min at 90 °C and for 60 min at 37 °C.

### Plasmid construction

To obtain plasmids for the production of *e*dsRNAs or control dsRNAs, a modified pUC18 (Thermo Scientific, Waltham, MA, #SD0051) containing two opposite T7 promoters flanking two BpiI (BbsI) restriction sites was first generated by inserting double-stranded oligonucleotides between the PstI and BamHI sites (Thermo Scientific, #ER0612 and #ER0051) of the vector. Prior to ligation using T4 DNA Ligase (Thermo Scientific, #EL0011), the oligonucleotides were phosphorylated by T4 Polynucleotide Kinase (Thermo Scientific, #EK0031). The plasmids containing the cDNAs of CMV *e*dsRNAs, consisting of several CMV *e*siRNA sequences of 21 or 22 nt length, were then generated by inserting double-stranded oligonucleotides into the BpiI-digested (Thermo Scientific, #ER1011) modified pUC18. The plasmids containing the cDNAs of the control RNAs dsCMV and dsGFP were generated by inserting respective PCR products, amplified with Phusion High-Fidelity DNA Polymerase (Thermo Scientific, #F530L) from plasmids pFny209 (see below) or pGFP-C1 (Clontech laboratories, Mountain View, CA), into the BpiI-digested modified pUC18. The resulting plasmids were used to generate PCR products, which served as templates for separate transcription of the two dsRNA strands.

### *In vitro* transcription

*Nicotiana benthamiana AGO1L* and *AGO2* mRNAs were transcribed *in vitro* from SmiI (SwaI)-linearized (Thermo Scientific, #ER1241) plasmids containing the corresponding open reading frames in a modified pSP64-poly(A) vector (Promega, Madison, WI) with the respective additional restriction site downstream of the poly(A) sequence (55,56). Transcription was performed in the presence of monomethylated cap analog m^7^GP_3_G (Jena Biosciences, Jena, Germany, #NU-852-5) using SP6 RNA Polymerase (Thermo Scientific, #EP0131). Firefly luciferase mRNA was produced by SP6 RNA Polymerase from plasmid pSP-luc(+) (Promega) linearized with XhoI (Thermo Scientific, #ER0692). CMV genomic RNAs (strain Fny) were synthesized with 5’ monomethylated cap analog by T7 RNA Polymerase (Agilent Technologies, Santa Clara, CA, #600123) from PCR product templates (see **Supplementary Table S1** for primer sequences). The template DNAs were amplified with Phusion High-Fidelity DNA Polymerase from plasmids pFny109, pFny209, and pFny309 (59) containing the respective cDNA sequences (kindly provided by Prof. Fernando García-Arenal Rodríguez, Universidad Politécnica de Madrid and Prof. John Carr, University of Cambridge). To generate double-stranded CMV RNA 2, non-capped antisense transcripts were synthesized from PCR products containing the T7 promoter in opposite orientation and subsequently annealed with non-capped sense transcripts. To achieve full complementarity also with the 5’-end of the antisense RNA (starting with guanosine due to the T7 promoter), the 3’-end of the sense RNA used for dsRNA production was changed from the original adenosine to cytidine. The two strands that constitute the *e*dsRNAs or the respective double-stranded control RNAs (dsCMV and dsGFP) were transcribed separately from individual PCR product templates that contained the T7 promoter in opposite orientations. Radioactive labeling of RNAs was performed by *in vitro* transcription in the presence of 0.2 µCi/µl [α-^32^P]-CTP (3,000 Ci/mmol, Hartmann Analytic, Braunschweig, Germany, #FP-209). Transcripts were treated with RNase-free DNase I (Roche Diagnostics, Mannheim, Germany, #04716728001) and purified by phenol/chloroform extraction and ethanol precipitation or by using the Nucleospin RNA Mini Kit (Macherey-Nagel, Düren, Germany, #740955.50).

### Preparation of double-stranded RNA

Annealing of RNAs was performed by mixing equimolar amounts of both single-stranded transcripts in STE buffer (10 mM Tris/HCl, pH 8.0, 100 mM NaCl, 1 mM EDTA), heating for 2 min at 94 °C and decreasing the temperature to 25 °C within 30 min.

### *e*NA screen

**Generation and characterization of siRNA pool** (see scheme in **Figure 1A**). 2.5 µg of the target-dsRNA were incubated for 2 h at 25 °C in a 100 µl reaction containing 50% (v/v) BYL, supplemented with 0.75 mM ATP, 0.1 mM GTP, 25 mM creatine phosphate, 80 μM spermine, 0.2 mg/ml Creatine Kinase (Roche Diagnostics, #10127566001) and 40% (v/v) TR buffer (30 mM Hepes–KOH, pH 7.4, 100 mM KOAc, 1.8 mM MgOAc, 2 mM DTT, cOmplete EDTA-free protease inhibitor cocktail (Roche Diagnostics, #05056489001)). RNA was isolated from the reaction by treatment with 1 µg/µl Proteinase K (Thermo Scientific, #EO0491) in the presence of 0.5% (w/v) SDS for 30 min at 37 °C, followed by phenol/chloroform extraction and ethanol precipitation. Purified RNAs from three reactions using different BYL preparations were combined and analyzed by RNA-Seq. Data were obtained from three independent experiments.

**Figure 1.**
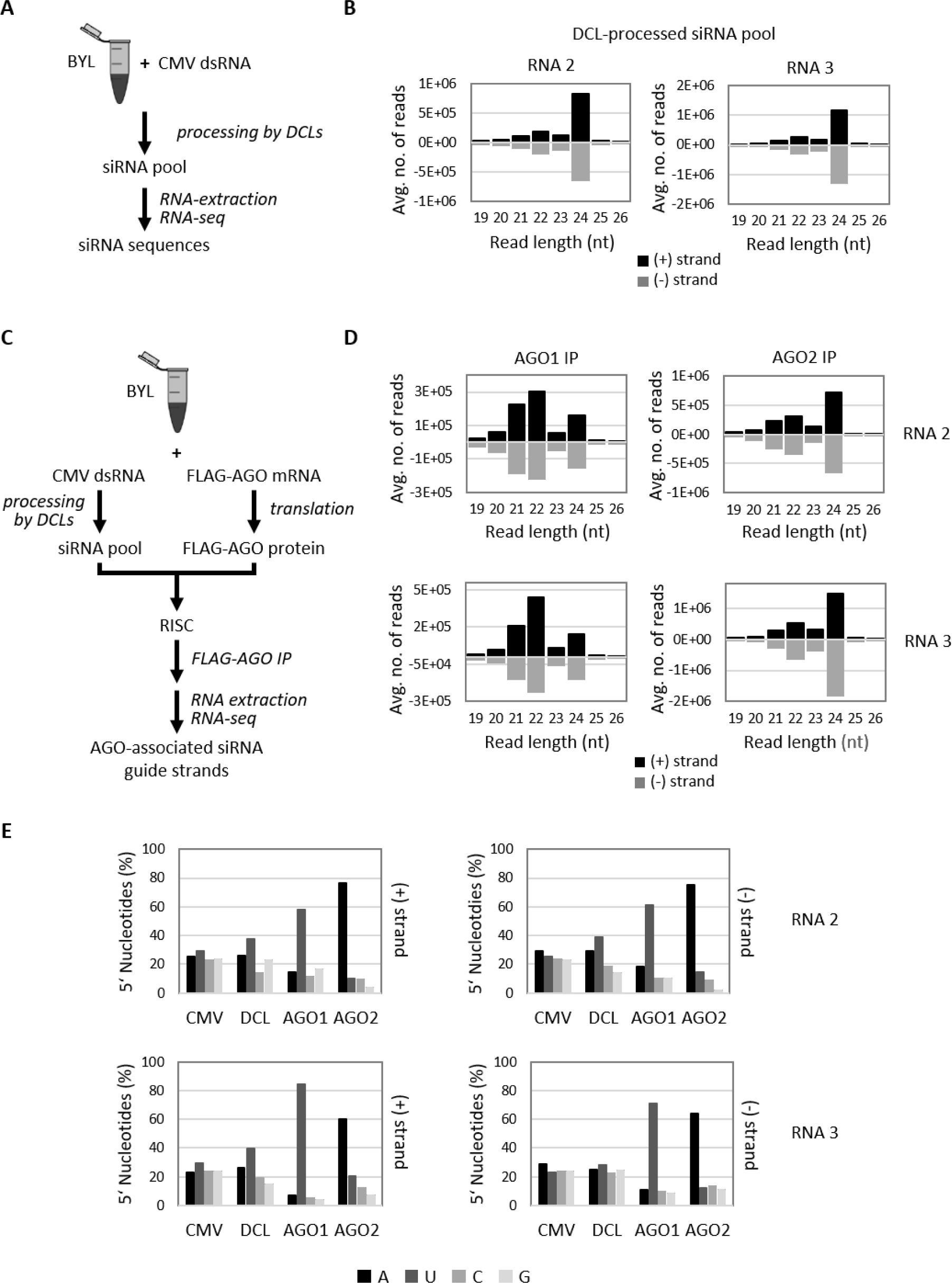
Identification and characterization of *e*siRNA candidates from CMV Fny RNAs 2 and 3 (*e*NA screens, steps 1 and 2). **(A)** Schematic of the siRNA pool generation and characterization procedure (see text for details). **(B)** Size distribution of siRNAs generated by the BYL-endogenous DCLs from dsRNA versions of CMV RNAs 2 and 3. Bars above and below the axis represent siRNAs derived from viral (+) and (-) strand RNA, respectively. Data represent the mean of three experiments. **(C)** Schematic representation of the procedure to identify AGO-bound siRNAs (see text for details). **(D)** Size distribution of CMV siRNAs isolated from AGO immunoprecipitations (IP). Data represent the mean of three experiments, except for AGO1 IP with CMV RNA 2 siRNAs (two experiments). **(E)** Relative abundance of the respective 5’-terminal nucleotides of the AGO1- and AGO2-associated 21 nt siRNA guide strands. The abundance was compared with the nucleotide compositions of the respective CMV RNAs (CMV), and the relative abundance of the 5’-terminal nucleotides of all sequenced 21 nt CMV siRNAs generated in BYL by endogenous DCLs (DCL; see **A** and **B**). Identified *e*siRNA candidates from CMV RNA 2 and CMV RNA 3 are listed in **Tables 1** and **2**, respectively.

**Characterization of AGO-bound siRNAs** (see scheme in **Figure 1C**). To generate siRNA-programmed AGO/RISC *in vitro,* 5 pmol *NbAGO1L* mRNA (56) or *NbAGO2* mRNA (60) were translated in the presence of 2.5 µg CMV dsRNA in a 100 µl reaction containing 50% (v/v) BYL under the above-described conditions. Both AGO proteins were produced with an N-terminal FLAG-tag. Samples were mixed with an equal volume of immunoprecipitation buffer (IPB, 20 mM Hepes/KOH, pH 7.6, 150 mM NaCl, 0.5% (v/v) NP-40, 1 mM DTT) containing 20 µl Anti-FLAG M2 affinity gel (Sigma-Aldrich, St. Louis, MO, #A2220). Following overnight incubation at 4°C with gentle agitation, the resin was washed three times with IPB, once with IPB containing 300 mM NaCl and a final time with IPB. RNA was isolated using proteinase K treatment, phenol/chloroform extraction and ethanol precipitation. Purified RNAs from three reactions using different BYL preparations were combined and analyzed by RNA-Seq. Data were obtained from three different experiments, except for the AGO 1 immunoprecipitation with CMV RNA 2-derived siRNAs (two experiments).

***In vitro* slicer (cleavage) assay** (see scheme in **Supplementary Figure 1C**). AGO/RISC programmed with a specific siRNA were generated as described above in a 20 µl reaction containing 50% (v/v) BYL, 0.5 pmol *NbAGO1L* or *NbAGO2* mRNA, 100 nM CMV siRNA duplex or 200 ng *e*dsRNA. After 2.5 h at 25 °C, 2 µg of firefly luciferase mRNA (serving as competitor RNA for non-specific acting RNases in BYL) and the ^32^P-labeled target RNA (10 fmol) were added, and the cleavage reaction performed for 15 min. Total RNA was isolated by treating the reaction with 20 µg Proteinase K in the presence of 0.5% SDS for 30 min at 37 °C, followed by chloroform extraction and ethanol precipitation. RNAs were separated on 1.5% denaturing agarose gels, ^32^P-labeled target RNAs and cleavage products were visualized by phosphor-imaging. To determine cleavage efficiencies of *e*siRNA candidates, the band intensities of the original, uncleaved transcripts were quantified with ImageQuant TL (Cytiva), data were obtained from two different experiments.

### RNA-Seq and bioinformatic analysis

RNA-seq was performed with an Illumina NextSeq 550 in the Core Unit DNA Technologies at the University of Leipzig (Germany), using the NEXTFLEX® Small RNA-Seq Kit v3 (Bioo Scientific, Austin, TX, #5132-05) to prepare the small RNA libraries. Adapter sequences were clipped from the raw reads using cutadapt 3.2 (61) with parameters specifying a maximum allowed error rate of 0.2 when matching adapters, a minimum overlap of 5 nt between adapter sequence and read, and a quality cutoff of 20 on the Phred-score from either end of the read, filtering for a minimum length of 15 nt of the adapter clipped reads (-e 0.2 -O 5 -q 20,20 -m 15). Since sequenced fragments were ligated with 4 nt random sequences at both ends prior to library preparation, additional 4 bases were cut from both ends of the adapter-clipped reads using cutadapt, again filtering for a minimum read length of 15 nt. Reads were mapped to the respective CVM RNAs using bowtie 1.3.0 (62). To this end, a bowtie index was generated including the chromosome and scaffold sequences of the *N. tabacum* Nitab 4.5 genome sequence (63) available from https://solgenomics.net/ftp/genomes/Nicotiana_tabacum/edwards_et_al_2017/assembly/ to avoid false-positive mappings of fragments originating from the cytoplasmic extract to the CMV RNAs. The bowtie command line was specified with additional parameters (--best --all --strata --tryhard -n 3) to achieve high mapping sensitivity. Reads mapping to the respective CMV RNA were extracted using samtools view 1.11 (64) for further processing, while reads mapping to the Nitab 4.5 sequences were discarded. Using custom Java and R scripts applied to the mapping result for each library, 5’-ends of reads mapping to the respective CMV RNA were counted position-wise, separately for each mapping strand and separately for each length fraction between 19 and 26 nt. Resulting count values were further used for determining the length distribution of mapped reads, the preference for 5’-nucleotides (based on the CMV RNA sequence), position-specific read counts and siRNA-specific read counts. Log2-fold changes between AGO1/AGO2 immunoprecipitation samples and DCL-processed siRNA pools were determined based on average count values, normalized to total library size per length fraction and treatment. Due to substantially different library sizes of replicates, normalization was performed prior to averaging in case of CVM RNA 3.

### RNA inoculation/CMV challenge of plants

Mechanical co-inoculation of siRNAs and CMV genomic RNAs was performed with 4-5 week-old *N. benthamiana* plants obtained from the Leibniz Institute of Plant Biochemistry (IPB) in Halle (Saale), Germany. Prior to the application of the RNAs, the upper surface of the third and fourth leaf was dusted with carborundum powder (silicon carbide, Sigma-Aldrich, #37,809-7). 5 µl solution containing *in vitro* transcribed CMV RNAs 1, 2, and 3 (20 fmol each) and siRNA (150 pmol, if not indicated differently) or dsRNA (70 pmol, if not indicated otherwise) were mixed with an equal volume of inoculation buffer (30 mM K_2_HPO_4_, pH 9.2, 50 mM glycine) and 2.5 µl were rubbed onto each leaf half using a pipette tip. Treated leaves were rinsed with water and the plants were grown in a chamber (CLF Plant Climatics, Wertingen, Germany) for 14 h at 23 °C, 90-100 µmolm^-2^s^-1^ light (at shelf level) and for 10 h at 21 °C in the dark. The plants were monitored daily for 4-5 weeks for the development of symptoms. Finally, leaf material was collected from representative symptomatic and asymptomatic plants and total RNA was isolated using TRIzol reagent (Thermo Scientific, #15596026) according to the manufacturer’s instructions. Data were obtained from at least two different experiments (see figures for number of plants), except for the control experiment with the single-stranded RNAs constituting the CMV *e*dsRNA (**Supplementary Figure 7B**).

## Results

### Identification of *e*siRNAs targeting CMV RNAs 2 and 3

CMV RNAs 2 and 3 were selected as targets for our antiviral approach because these genome segments each encode two proteins essential for the infectious viral life cycle. CMV RNA 2 encodes the viral RNA-dependent RNA polymerase 2a and, *via* the subgenomic RNA 4A, the viral suppressor of RNA silencing (VSR) 2b. CMV RNA 3 encodes the movement protein (MP) 3a and, *via* the subgenomic RNA 4, the viral capsid protein CP (**Supplementary Figure 1B**).

To obtain *e*siRNAs, we adapted a previously developed protocol for the *e*NA screen (32) (see Materials and Methods for details), which consists of three steps that are summarized here and explained in detail in the following sections. (i) Delivered as dsRNA, the target RNA of choice was exposed to BYL for processing by endogenous DCLs to produce a pool of siRNAs. Next-generation sequencing (RNA-seq) monitored the entire siRNA population (scheme in **Figure 1A**). (ii) In BYL containing such a siRNA pool, RISCs were reconstituted with an AGO protein of choice. AGO was translated *in vitro* from a cognate (*in vitro* transcribed) mRNA. Formed AGO/RISC were immunoprecipitated, and the bound siRNA strands were identified by RNA-seq (scheme in **Figure 1B**). The RNA-seq data obtained in step (i) were then used for comparison to define enrichment in the AGO/RISC. (iii) From the siRNAs detected in step (ii), those siRNAs that induced efficient slicing of the target RNA were finally identified by BYL-supported slicer assays that test for endonucleolytic cleavage of the target RNA (see **Supplementary Figure 1C** for assay scheme).

Applying this approach to CMV, full-length sense and antisense transcripts were generated from cDNA clones of RNAs 2 and 3 of the Fny strain (59) and hybridized (see Materials and Methods). The resulting dsRNAs were processed from BYL-endogenous DCLs, and the generation of small RNAs was determined qualitatively by gel electrophoresis and quantitatively by RNA-seq (**Supplementary Figure 2A** and B; **Figure 1B**). In close agreement with previous *e*NA screens (32), siRNAs with lengths of 21 nt to 24 nt were detected, with the ratio of siRNAs generated from the positive- and negative-orientated strands of the dsRNAs being balanced. Consistent with previous observations, siRNAs of 24 nt were found to be the predominant type (**Figure 1B**) suggesting that DCL3, which processes dsRNA into 24 nt long siRNAs, is more abundant or more active in BYL than DCL2 and DCL4. Since 21 nt siRNAs were of particular interest due to their documented antiviral activity, we analyzed the average frequency with which a 21 nt siRNA was processed at a specific position of the respective dsRNA versions of genomic CMV RNAs 2 and 3. The data showed that DCL processing occurred along the entire length of the RNAs, with slightly higher frequencies at certain sites (**Supplementary Figure 2B**). Analogous to data previously obtained with dsRNA of the genome of Tomato Bushy Stunt Virus (TBSV), which is not related to CMV, this behavior of DCL4, the enzyme most likely involved here, could not be explained by any general rule, except for a certain preference for cleavage at the RNA termini and at GC-rich sequence motifs (32,65) (data not shown).

To identify those virus-derived siRNAs whose guide strands are preferentially bound by the AGO1 and AGO2 proteins, we repeated the generation of the CMV RNA-derived siRNA pools and then exposed the siRNAs to AGO1 (AGO1L variant from *N. benthamiana*; (56)) or AGO2 (from *N. benthamiana*; (60)), which were translated *in vitro* in BYL. The AGOs were produced with an N-terminal FLAG tag, which allowed immunoprecipitation (IP) of the proteins including the bound siRNAs. Successful translation and IP of the FLAG-AGO proteins was verified as well as co-precipitation of DCL-processed siRNAs with the corresponding FLAG-AGO (**Supplementary Figure 2C** and D). Subsequent RNA-seq of the precipitated siRNAs revealed a massive enrichment of 21 nt and 22 nt siRNAs compared to 24 nt siRNAs, especially with AGO1, but also AGO2 (compare **Figure 1D** and **1B**). Consistent with the known preferences of AGO proteins for binding siRNA guide strands, the vast majority of siRNAs bound by AGO1 had a 5’-terminal uridine, whereas siRNAs bound by AGO2 predominantly had a 5’-terminal adenosine (**Figure 1E; Supplementary Table 2**). Further analyses of the RNA-seq data focused on 21 nt siRNAs that were significantly enriched in the respective AGO/RISC and whose guide strands had negative strand polarity, *i.e.,* they were complementary to the CMV genomic RNAs. Thus, siRNAs were selected that were highly enriched in the precipitated AGO/RISC compared to the DCL-processed siRNA pool, or siRNAs that were only detected in the IP samples but not in the original siRNA pool: both criteria indicated a high affinity of the siRNA to the corresponding AGO protein. In the case of CMV RNA 2, these were 12 AGO1- and 14 AGO2-bound siRNAs; in the case of CMV RNA 3, these were 11 AGO1- and 13 AGO2-bound siRNAs, which were considered for further study (**Supplementary Table 2**; **Tables 1** and **2**). For unknown reasons, some of the candidates contained a 5’-terminal nucleotide that did not match AGO1’s and AGO2’s major preferences (**Supplementary Table 2**). Tests with examples of these siRNAs (*e.g.* siR1844) showed that they were not functional (**Figure 2**, and data not shown). Therefore, from the outset, siRNAs were excluded from further analysis if their 5’-nucleotides did not match the AGO-specific preferences.

**Figure 2.**
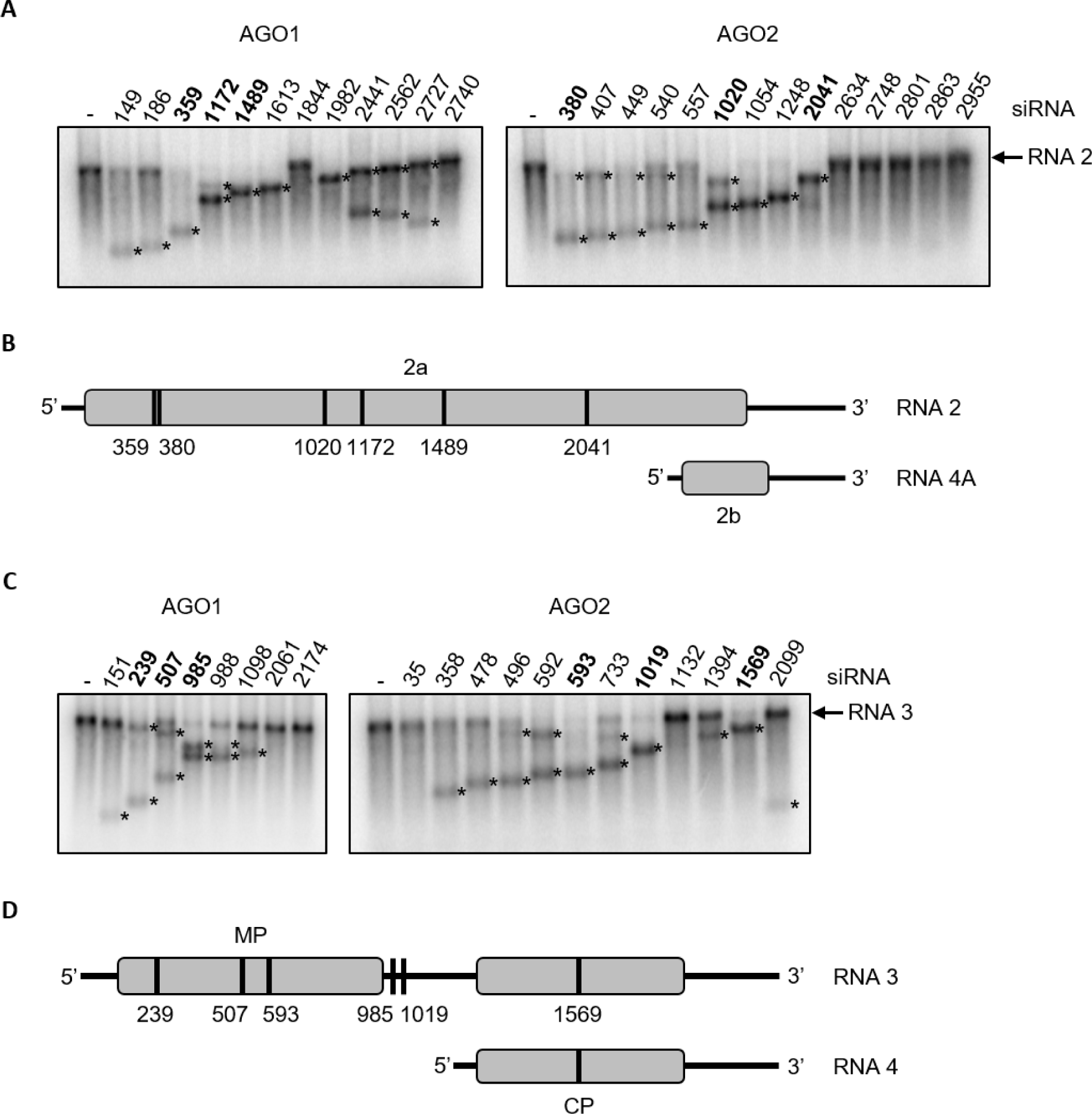
RNA silencing activity of *e*siRNA candidates *in vitro* (*e*NA screens step 3). Slicer assays with the *e*siRNA candidates identified in step 2 of the *e*NA screen and CMV RNAs 2 or 3 (Figure 1) were performed as shown schematically in **Supplementary Figure 1C** and described in the text. The numbers of the *e*siRNA candidates correspond to the designations given in the text, tables and figures below. Asterisks (*) denote the cleavage products. Determined slicing efficiencies are summarized in **Tables 1** and **2**. **(A)** Representative slicer assays performed with *e*siRNA candidates, CMV RNA 2 and AGO1/RISC or AGO2/RISC. Six *e*siRNA candidates (three active with AGO1, three active with AGO2) that were further investigated in the following are shown in bold. **(B)** Schematic representation of the binding sites of these siRNAs on CMV RNA 2. **(C)** Representative slicer assays performed with *e*siRNA candidates, CMV RNA 3 and AGO1/RISC or AGO2/RISC. *e*siRNA candidates that were further investigated are shown in bold. **(D)** Binding sites of these siRNAs on CMV RNA 3.

When tested *in vitro* for slicer/target cleavage activity, most, but not all, of the selected siRNAs mediated substantial cleavage of the CMV RNAs 2 and 3 (**Figure 2**). In fact, based on data from various *in vitro* screening studies, the majority of these siRNAs met the arbitrary definition of an “*e*siRNA candidate”: accordingly, a candidate *e*siRNA was defined here as one that induces hydrolysis of more than 25% of a target RNA in a standardized *in vitro* slicer assay with appropriate AGO/RISC compared to a control reaction without siRNA (**Figure 2**; see also Materials and Methods). For example, based on their cleavage activity, siR359, siR1172 and siR1489 were identified as *e*siRNA candidates with AGO1 on CMV RNA 2 (candidates are named with “siR” and the position of the viral positive-strand RNA to which the 5’-nucleotide of the siRNA guide strand is complementary). Similarly, siR380, siR1020 and siR2041 were identified as highly cleavage-active siRNA candidates with AGO2/RISC (**Table 1**). Examples of *e*siRNA candidates that acted with AGO1 on CMV RNA 3 were siR239, siR507, and siR985; with AGO2 they were siR593, siR1019 and siR1569 (**Table 2**). In general, however, both the total number and the cleavage efficiencies of the *e*siRNA candidates identified for CMV RNA 3 were significantly lower than for CMV RNA 2 (**Figure 2**; **Tables 1** and **2)**.

**Table 1.**
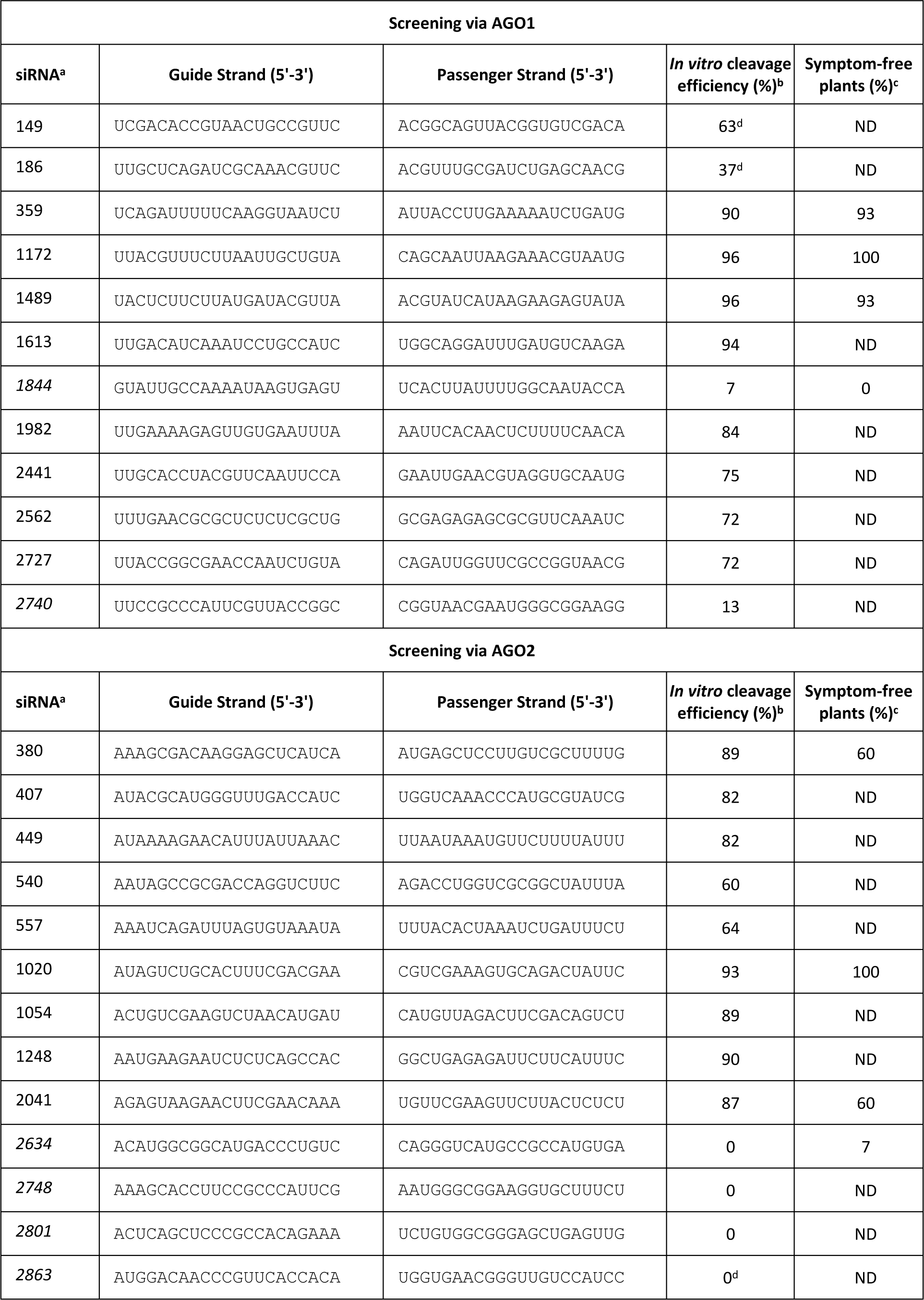
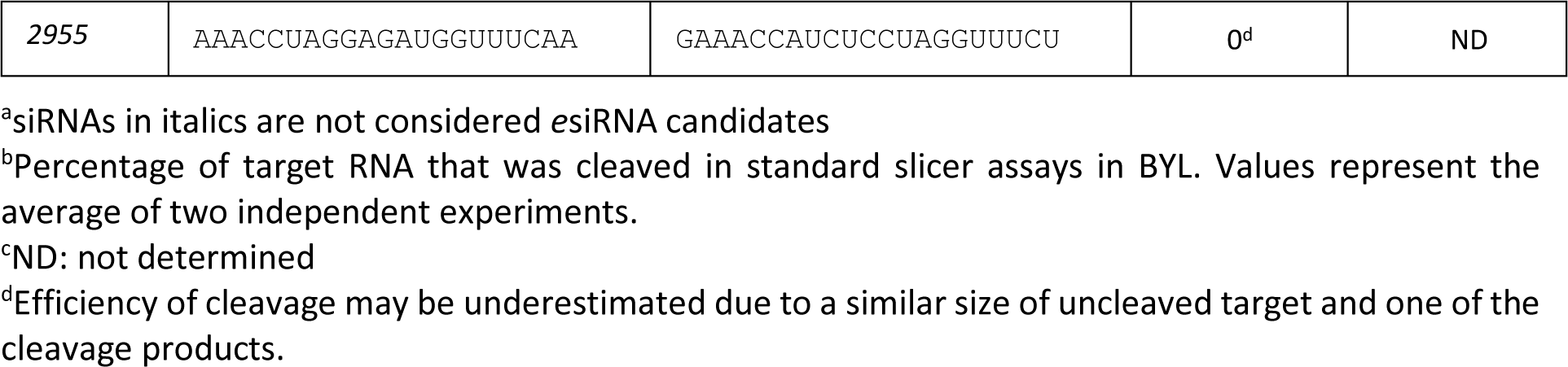
*e*siRNA candidates targeting CMV RNA 2.

**Table 2.**
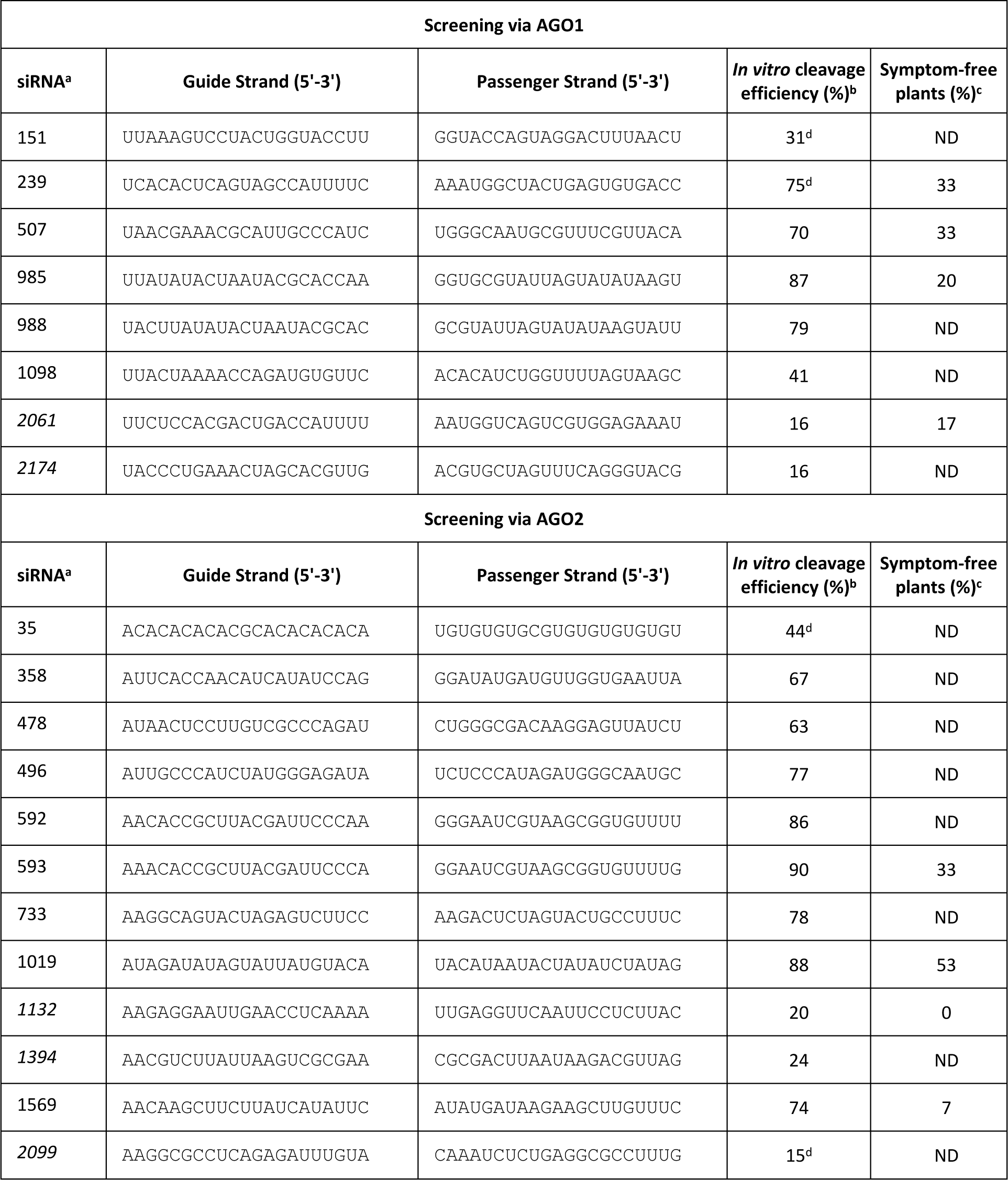
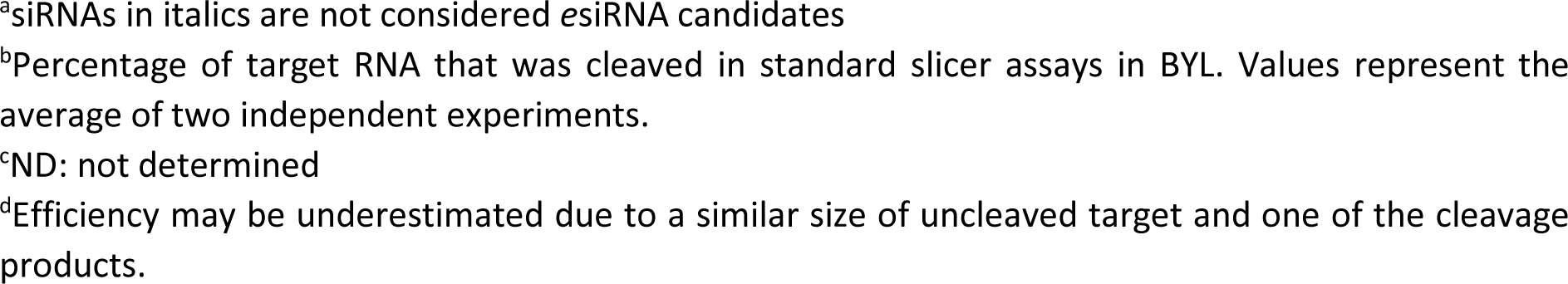
*e*siRNA candidates targeting CMV RNA 3.

Subsequently, the antiviral (protective) activity of some of the identified *e*siRNA candidates was tested *in vivo*. For this purpose, *N. benthamiana* plants (n=12-15) were treated with carborundum using a standard protocol (“rub-inoculation”) and 150 pmol (∼1 µg) of the (synthesized) siRNA to be tested. At the same time, 20 fmol of each of the CMV genomic RNAs 1-3, prepared by *in vitro* transcription from full-length cDNAs, were inoculated per plant. It is important to note that this amount of CMV Fny RNAs, when inoculated into *N. benthamiana* plants without the addition of antiviral siRNAs was previously confirmed to result in infection of 100% of the plants with significant symptom development and pathogenesis, including deformed leaves and severe dwarfism. This means that the plants were subjected to what we refer to as a “maximally successful challenge” by the virus. First, the 21 nt long siRs 359, 380, 1020, 1172, 1489 and 2041 derived from CMV RNA 2 were used. As a negative control, we applied siR gf698, a non-specific siRNA-targeting green fluorescent protein (GFP) mRNA. In addition, we used two siRNAs, siR1844 and 2634, which showed no, or low, cleavage activity on the target RNA in the previous slicer assays (**Figure 2**; **Table 1**). **Figure 3A** shows representative images of plants treated as described above and examined for symptom development at a maximum of 35 days post-inoculation (dpi); **Figure 3B** shows the overall progression over several independent infection/protection experiments (three independent experiments including 4 to 5 plants per treatment/per experiment). It was found that siR359 and siR1489 each provided 93% protection and siR1020 and siR1172 each provided 100% protection against CMV infection, indicating that all, or the vast majority, of the plants treated in this way remained symptom-free. siR380 and siR2041 provided 60% protection against CMV infection, meaning that 40% of these plants developed symptoms. The control siRNA and, importantly, the two siRNAs that were inactive in previous slicer assays provided little or no protection; since none, or very few (7%), of the plants treated in this way remained symptom-free (**Figure 3A** and **B**). These results show that the observed protective effect of siRs 359, 380, 1020, 1172, 1489 and 2041 was indeed due to RNA-silencing processes in the plant. In other words, we concluded that the protective effect was not triggered by single-stranded siRNA guide strands that might still be present as contamination in the siRNA preparations and could hybridize to the corresponding viral RNA strands during the inoculation process (see also below). Protection was also confirmed by the fact that genomic CMV RNA was no longer detectable by RT-PCR in plants identified as asymptomatic at 35 dpi (**Supplementary Figure 3**; all data summarized in **Table 1**).

**Figure 3:**
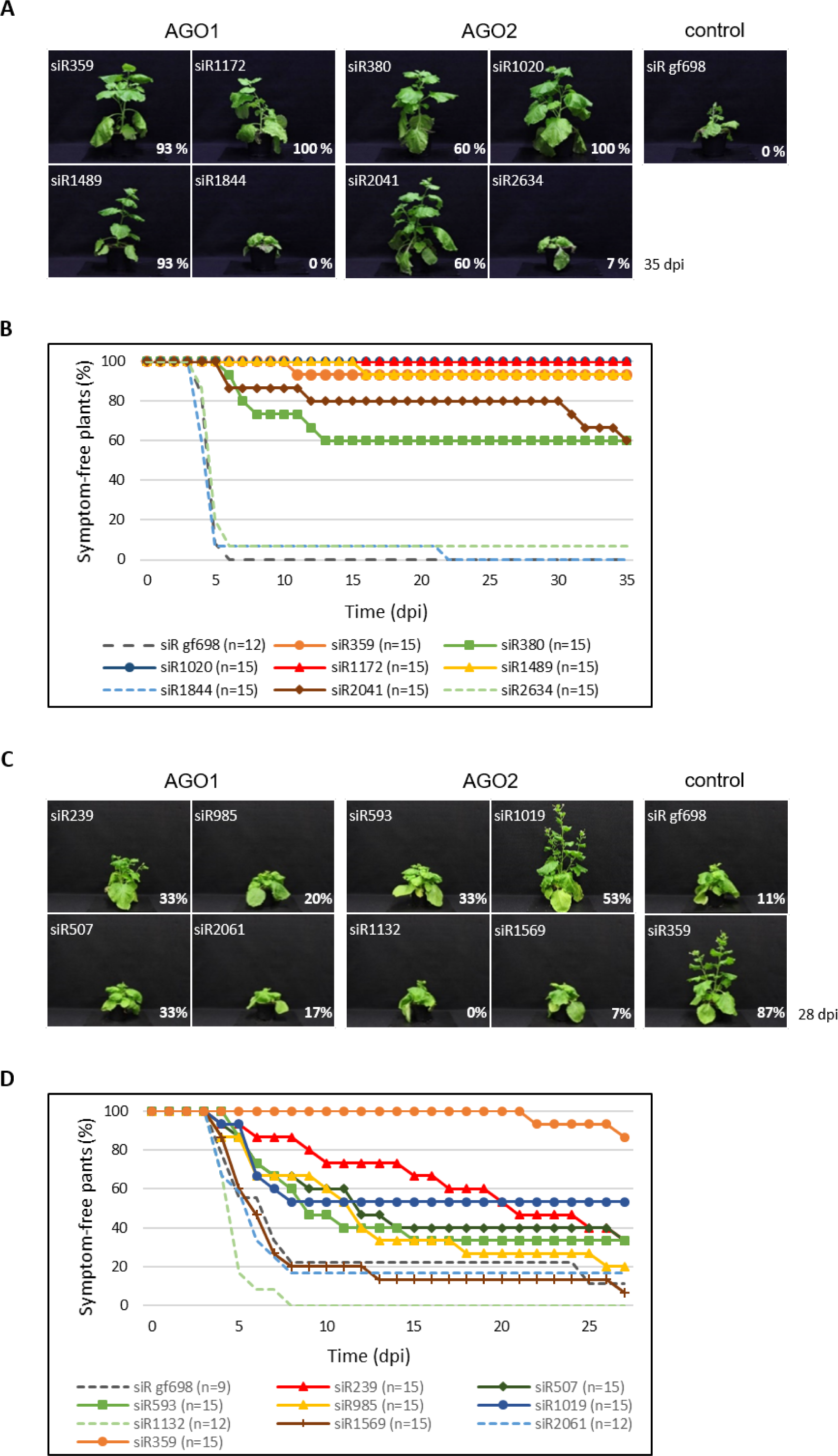
Identified *e*siRNAs protect plants efficiently against CMV infection. *N. benthamiana* plants were mechanically co-inoculated with the individual synthetic *e*siRNA candidates and with the genomic CMV RNAs 1, 2, and 3. The concentration of the viral RNAs was chosen to achieve a maximal successful challenge with the pathogen (see text). siRNA gf698 targeting GFP mRNA was used as a negative control. Plants were monitored for the appearance of CMV-specific symptoms for at least 28 dpi. **(A, C)** Representative plant images illustrate the differences between asymptomatic and symptomatic individuals at 35 dpi (RNA 2-targeting siRNAs) or at 28 dpi (RNA 3-targeting siRNAs). The percentage of asymptomatic remaining plants is given for each siRNA. **(B)** Percentage of asymptomatic plants over the entire course of the experiment with RNA 2-targeting siRNAs. Results are from three independent experiments, the total number of plants is indicated (n = 12-15). **(D)** Percentage of asymptomatic plants over the entire course of the experiment with RNA 3-targeting siRNAs. Results are from three independent experiments, the total number of plants is indicated (n = 9-15). An *e*siRNA directed against CMV RNA 2 (siR359) was used as an additional control.

The equivalent experiment with the candidates characterized from CMV RNA 3 showed that protection against CMV infection could also be achieved here. However, the level of protection was clearly lower than with the *e*siRNA candidates from CMV RNA 2 (**Figure 3C** and **D**; summarized in **Table 2**). This was expected, as most of these siRNAs directed against CMV RNA 3 showed a lower slicer activity *in vitro* (**Figure 2**; **Tables 1** and **2**). Note that an *e*siRNA directed against CMV RNA 2, siR359, was also tested in the experiments shown in **Figure 3C** and **D**, and it provided comparable protection to the infection experiments shown in **Figure 3A** and **B**. This demonstrates the high degree of reproducibility of these experiments.

Like 21 nt siRNAs, 22 nt siRNAs were previously shown to play an important role in establishing effective RNAi responses (15,66). Accordingly, we next wanted to know whether a comparable antiviral protection could be achieved with 22 nt versions of the previously characterized *e*siRNAs (guide strands extended by one nucleotide). In slicer assays, the 22 nt siR359, siR1172, siR1489, siR380, siR1020 and siR2041 were found to have similar or slightly lower activity in the respective AGO/RISC on the target RNA (**Supplementary Figure 4A**). Plant protection experiments with the 22 nt siR359 and siR1020 showed a similar tendency: *i.e.,* the protection levels were either similar (siR359) or reduced (siR1020) compared to those of the 21 nt *e*siRNAs (exemplarily shown in **Supplementary Figure 4B** and C).

Thus, from an undefined siRNA pool, *e*NA screening reliably characterized *e*siRNAs that were functional, *i.e.* highly effective in topical antiviral applications. 21 nt versions were more potent than 22 nt versions of these *e*siRNAs, and CMV RNA 2-derived *e*siRNAs were more potent than CMV RNA 3-derived *e*siRNAs.

“Multivalent” double-stranded RNAs, “*e*dsRNAs”, composed of the sequences of several *e*siRNAs, are highly effective in protecting plants against CMV infection

For crop protection applications, dsRNAs are considered a better choice than siRNAs, because they are significantly cheaper to produce (23) and more stable (67). As mentioned above, currently used antiviral dsRNAs consist only of complementary long, contiguous regions of viral genomic RNA. From these, few, if any, *e*siRNAs were produced during the DCL-mediated processing. In addition, many other non-functional or undesirable (*e.g.,* potentially “off-targeting”) siRNAs were produced (see also Discussion). After the successful identification of *e*siRNAs against CMV, the use of dsRNAs containing several *e*siRNA sequences seemed particularly promising; DCLs should generate the constituent *e*siRNAs, and the corresponding RISCs should then attack the target RNA at different a-sites to hydrolyze it with maximum efficiency. Therefore, engineering cDNAs to produce “*e*dsRNAs” that are effective against CMV was the next step in this study.

It was particularly important to ensure that the *e*siRNAs that compose the *e*dsRNA are actually produced by the DCLs, especially since the way in which plant DCLs enzymatically convert dsRNA substrates is incompletely characterized (see Introduction section). Previous and recent observations (68–70) support the hypothesis that DCLs should be active at both ends of a dsRNA and endonucleolytically hydrolyze the RNA from there, possibly in a phased pattern. This was the basis for the construction of cDNAs that allowed the transcription of complementary strands into *e*dsRNAs. These cDNAs contained the sequences of six of the previously characterized 21 nt or 22 nt *e*siRNAs 1172, 1489, 359 and 1020, 2041, 380, respectively, with the sequences of the guide strands of three *e*siRNAs arranged in opposite directions initiating from the respective 5’-end (see **Figure 4A**; **Supplementary Figure 5**). To obtain intact double strands, the 2 nt overhangs of the passenger strands of *e*siRNAs 359, 380 and 2041 were modified to match the corresponding complementary bases.

**Figure 4.**
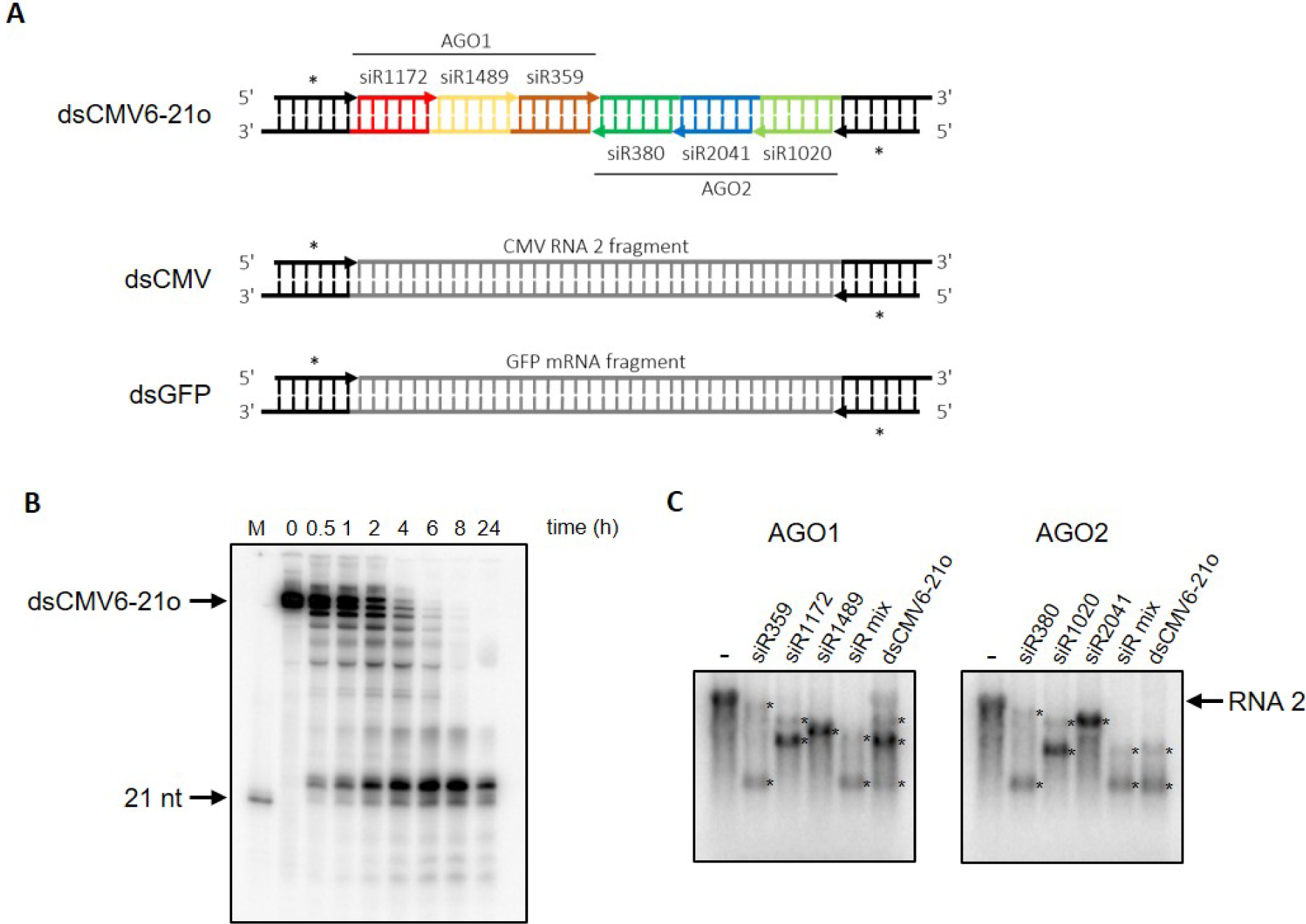
CMV *e*dsRNAs: composition, processing and *in vitro* silencing activity. **(A)** Exemplary *e*dsRNA and control dsRNAs. Top: Composition of an *e*dsRNA generated from two complementary RNA transcripts. The *e*dsRNA dsCMV6-21o contains a 21 nt long pseudo-siRNA at each end (symbolized by asterisks (*)) and six 21 nt long *e*siRNA sequences. These *e*siRNAs have been shown to be effective against CMV RNA 2 *in vitro* and to be antivirally protective *in planta* (numbering according to Figure 2 and **Table 1**). Guide strands (gs) are shown as arrows pointing in the 5’-3’ direction. The AGO1-specific gs are located on one RNA strand, the AGO2-specific gs are located on the other RNA strand. The example RNA shown here has 2 nt 3’ overhangs (dsCMV6-21o); however, *e*dsRNAs with blunt ends (dsCMV6-21) were also generated and tested. In addition, *e*dsRNAs containing the sequences of 22 nt pseudo-siRNAs at the ends and six 22 nt *e*siRNAs (dsCMV6-22o; dsCMV6-22) were generated and tested (see text). Middle: Control RNA dsCMV consists of pseudo-siRNA sequences at the ends and a 126 nt-long fragment of a double-stranded version of CMV RNA 2 (corresponding to a length of six 21 nt-long siRNAs). By chance, the dsCMV also contains the sequences of two siRNAs that were identified as *e*siRNAs in the screen against CMV RNA 2. Bottom: Control RNA dsGFP also consists of pseudo-siRNA sequences at the ends and a 126 nt long fragment of a double-stranded version of GFP mRNA. The exact sequences of the dsRNAs shown are given in **Supplementary Figure 5**. (**B**) Processing of an *e*dsRNA by DCLs *in vitro*. Labeled dsCMV6-21o was added to BYL and DCL-mediated processing analyzed over 24 h (see Materials and Methods) (M = 21 nt siRNA as marker). (**C**) Slicer assays with individual AGO1- and AGO2-specific *e*siRNAs from CMV RNA 2 and with the analogous *e*siRNAs generated from an *e*dsRNA in BYL. AGO1- or AGO2/RISC were reconstituted with individual *e*siRNAs, with an appropriate mix of these *e*siRNAs or with *e*siRNAs processed from the *e*dsRNA dsCMV6-21o in BYL by the DCLs present there. Endonucleolytic hydrolysis of labeled CMV RNA 2 target was detected by gel electrophoresis and autoradiography. Asterisks (*) indicate the generated cleavage products.

As an additional feature of *e*dRNA-encoding cDNAs, we have introduced 21 or 22 nt-long, so-called “pseudo-siRNA sequences” (**Figure 4A**). These served two purposes: at the cDNA level, these elements contained consensus sequences (+1 to +6) that promote T7 RNA polymerase-mediated transcription; otherwise, the nucleotide composition was random. Both *e*dsRNA strands could be generated accordingly by *in vitro* transcription. At the RNA level, we hypothesized that the presence of the appropriate length pseudo-siRNA sequences should support the proper processing of the downstream *e*siRNA components. Assuming that DCL4, like other DCLs and Dicers, acts in a processive manner using the 5’-counting rule (6,68,71), the 21 nt-long *e*siRNAs should preferentially result from DCL4 activity on this substrate if the pseudo-siRNAs are 21 nt long. A similar scenario was expected for DCL2 and edsRNA substrates with terminal 22 nt pseudo-siRNA sequences. The cDNAs encoding the 21 or 22 nt pseudo-siRNAs and *e*siRNAs were used to generate PCR products that served as templates for separate *in vitro* transcription of the single-stranded components of the dsRNA. Subsequently, the transcripts were hybridized to obtain the dsRNA. By using different PCR primers, we generated dsRNAs with blunt ends (here referred to as dsCMV6-21 or dsCMV6-22; see also **Figure 4A** and **Supplementary Figure 5** for the sequence composition), or dsRNAs with 2 nt-long 3’-overhangs (here referred to as dsCMV6-21o or dsCMV6-22o). The latter variants were developed based on reports suggesting that DCLs on dsRNAs with overhangs show more precise processing (68,72).

First, we tested the DCL-mediated processing of the constructed *e*dsRNAs. To do this, we again used the endogenous DCL activities in BYL; in analogy to the experimental procedure shown in **Figure 1A**, labeled *e*dsRNAs were added to BYL and the processing to siRNAs was monitored over a period of 24 hours. For example, with dsCMV6-21o, we obtained a processing pattern indicating the generation of 21 nt siRNAs. However, in line with the fact that BYL contains not only active DCL4 but also DCL3, significant amounts of 24 nt-long siRNAs were also generated (**Figure 4B**).

Next, we tested the *e*dsRNAs in a slicer assay. Analogous to the previous procedure (**Figure 4B**), the *e*dsRNAs were processed in BYL by the endogenous DCLs. AGO1/RISC or AGO2/RISC were reconstituted with the resulting siRNAs and slicing of CMV RNA 2 target was analyzed (**Supplementary Figure 1C**). The assays with the *e*dsRNAs were performed side-by-side with the individual *e*siRNAs or with a mixture of the respective *e*siRNAs. As shown in **Figure 4C** with dsCMV6-21o, efficient hydrolysis of CMV RNA 2 occurred in all cases, *i.e.,* the target RNA was almost completely degraded to the corresponding cleavage products. Furthermore, efficient cleavage was obtained regardless of whether the *e*dsRNAs were composed of 21 nt or 22 nt *e*siRNA sequences and whether they contained a 2 nt overhang or not (**Supplementary Figure 6**). Systematic testing revealed that approximately 0.2 pmol (approximately 25 ng) of *e*dsRNA was sufficient to induce complete hydrolysis of 20 fmol of CMV RNA-2 target in a standard slicer assay (not shown). Taken together, these data demonstrate that the *e*dsRNA constructs used are processed by DCLs under the conditions of the *in vitro* BYL system, such that the resulting siRNAs induce AGO/RISC-mediated hydrolysis of the original viral target RNA with efficiencies similar to those of the individual *e*siRNAs.

To determine the extent to which the DCL-processed siRNAs corresponded to the original *e*dsRNA-constituting *e*siRNAs, we performed the following experiment. In line with the scheme shown in **Figure 1A**, the *e*dsRNAs dsCMV6-21 and dsCMV6-21o were exposed to the BYL-endogenous DCLs. After DCL-mediated processing, total RNA was extracted and small RNAs were sequenced using RNA-seq. Analysis of the size distribution of the siRNAs (**Figure 5A**) showed almost identical patterns for the two dsRNA constructs. Due to the dominant activity of DCL3 in BYL, and in agreement with previous observations (**Figures 1B** and **4B**), the predominant length of siRNA species processed from both *e*dsRNAs was 24 nt. However, in contrast to the previous data, the proportion of 21 nt siRNAs was significantly higher compared to 22 nt siRNAs (compare **Figure 5A** and **1B**). The most interesting results were obtained when we quantified the guide and passenger strands of the *e*dsRNA-forming siR1172, siR1489, siR359, siR380, siR2041, siR1020 and the pseudo-siRNAs at the termini: this revealed a significant dominance of the read abundances of these same siRNAs compared to the total number of 21 nt reads (total number of 21 nt siRNAs generated) (**Figure 5B** and **Supplementary Figure 7A**). Approximately 60% of all 21 nt reads corresponded to the guide and passenger strands of the *e*dsRNA-constituent *e*siRNAs and pseudo-siRNAs, respectively (**Table 3**). In most cases, the guide strands were more readily measurable, and the detected amounts differed between the individual *e*siRNAs, which was explained by different protection against RNases, biases in the RNA-seq procedure, but also by the DCL activities on these *e*dsRNAs. Furthermore, the data obtained indicate that the DCL4 molecules present in BYL do indeed process the *e*dsRNAs predominantly in a phased manner, and that an *e*dsRNA consisting of a 21 nt pseudo-siRNA and *e*siRNA sequences produces predominantly 21 nt *e*siRNAs, as expected. However, DCL4 was not the only active DCL and, accordingly, other siRNA species were also produced from the *e*dsRNAs. Since the results with the blunt- and 3’-overhang *e*dsRNAs were the same, we concluded that the nature of the *e*dsRNA termini has little effect on DCL-mediated processing in the plant extract (see Discussion).

**Figure 5.**
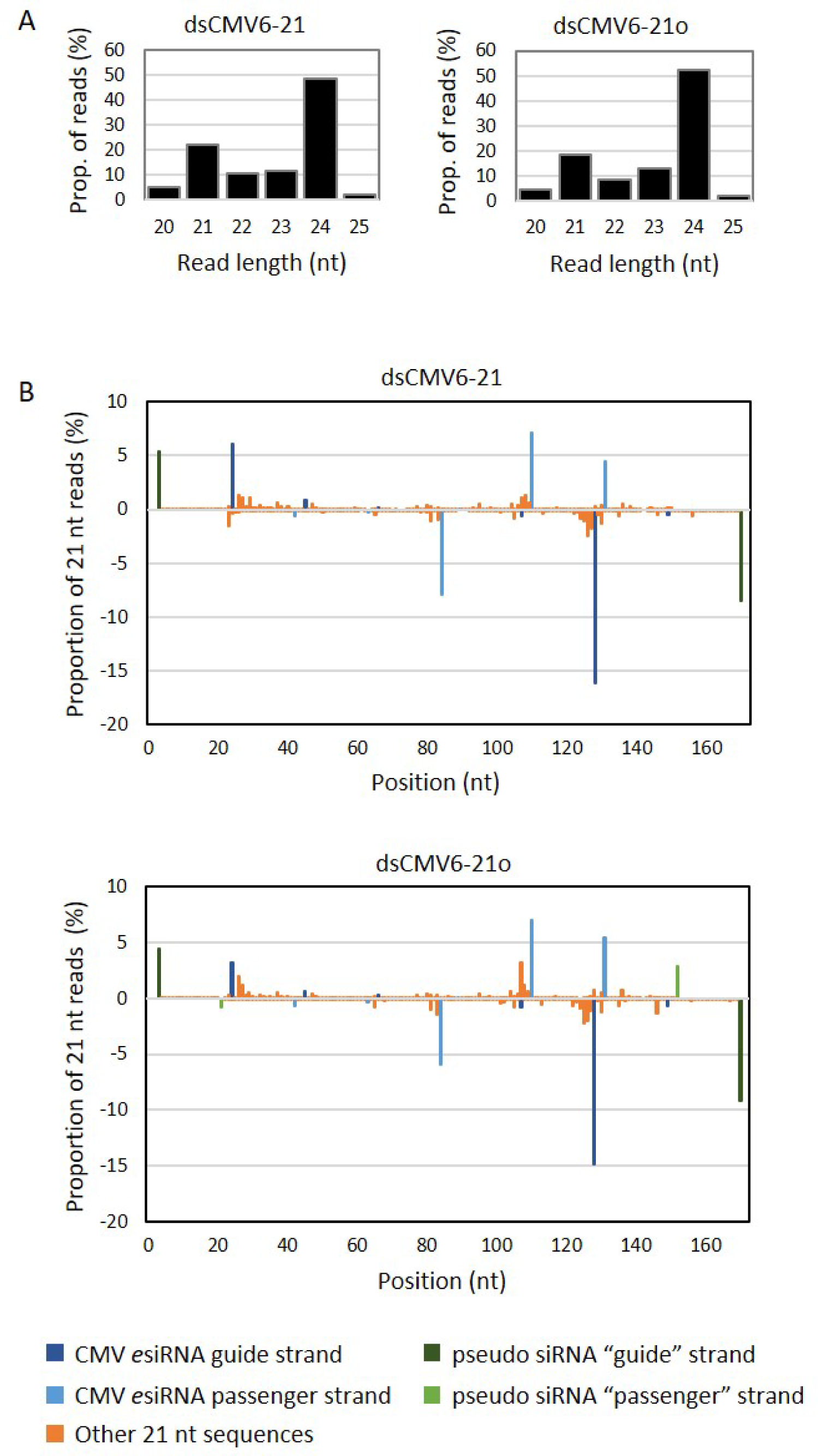
The *e*siRNA constituents are generated at high proportion from *e*dsRNA. dsCMV6-21 (blunt ends) and dsCMV6-21o (2 nt 3’ overhangs) were processed in BYL by the endogenous DCLs and the small RNA fraction analyzed by RNA-seq (see also scheme in Figure 1A). (**A**) Size distribution of the 20-25 nt reads mapping to the *e*dsRNA sequences. (**B**) Proportion of guide and passenger strand reads among all 21 nt reads that mapped to the *e*dsRNA sequences; the peaks indicate the position of the 5’-nucleotide of reads with respect to the *e*dsRNA. Peaks corresponding to the *e*dsRNAs contained pseudo-siRNA and *e*siRNA sequences are specifically colored (blue, green). In the case of dsCMV6-21 there are no mapping 21 nt reads for the “passenger” strands of the pseudo siRNAs, as the two 3’-nucleotides (position 20 and 21) are missing.

**Table 3.** Proportion of CMV-specific *e*siRNA strands among all 21 nt siRNAs that were processed from two types of *e*dsRNAs in BYL.

| siRNA | dsCMV6-21 |  | dsCMV6-21o |  |
| --- | --- | --- | --- | --- |
|  | Guide Strand (%) | Passenger Strand (%) | Guide Strand (%) | Passenger Strand (%) |
| pseudo 1<sup>a</sup> | 5.40 | - | 4.46 | 0.82 |
| 1172 | 6.09 | 0.65 | 3.19 | 0.66 |
| 1489 | 0.85 | 0.32 | 0.60 | 0.35 |
| 359 | 0.14 | 7.88 | 0.32 | 5.91 |
| 380 | 0.67 | 0.05 | 0.79 | 0.06 |
| 2041 | 16.18 | 7.18 | 14.91 | 6.96 |
| 1020 | 0.54 | 4.45 | 0.74 | 5.36 |
| pseudo 2<sup>a</sup> | 8.55 | - | 9.22 | 2.91 |
| total | 38.42 | 20.52 | 34.23 | 23.04 |

<sup>a</sup>Here, the strand with the same orientation as the adjacent siRNA guide strands was designated as “guide strand”, regardless of the probability of loading into AGO proteins.

The protective antiviral potential of *e*dsRNAs was evaluated in the final experiments of this study. To this end, we performed protection/infection experiments with *N. benthamiana* plants inoculated with 20 fmol of the three CMV genomic RNAs for a “maximally successful challenge” and approximately 70 pmol (8 µg) of dsCMV6-21o or dsCMV6-22o, respectively. As a control, we used a conventionally organized dsRNA, which we will refer to here as dsCMV (shown schematically in **Figure 4A**; for sequence composition, see **Supplementary Figure 5**). dsCMV consists of the previously described pseudo-siRNA sequences at the termini and a 126 nt-long continuous double-stranded portion of CMV RNA 2 (corresponding to nucleotide positions 501-626 of CMV RNA 2 and a length of six 21 nt-long siRNAs). Importantly, this double-stranded segment of CMV RNA 2 contained the sequences of two siRNAs, siR557 and siR540, which were identified as *e*siRNAs in the previous screening procedure (see above and **Supplementary Figure 5**). A double-stranded segment of GFP mRNA, dsGFP, served as a negative control: as in all other cases, this dsRNA corresponded to a length of six siRNAs and contained pseudo-siRNA sequences at the termini. In addition, dsGFP contained the sequence of the siR gf698 control siRNA that was used previously (**Supplementary Figure 5**). **Figure 6A** shows representative images of plants treated as described, and **Figure 6B** shows the overall course of several independent experiments performed with a representative number of plants and in which symptom development was monitored for 35 dpi.

**Figure 6.**
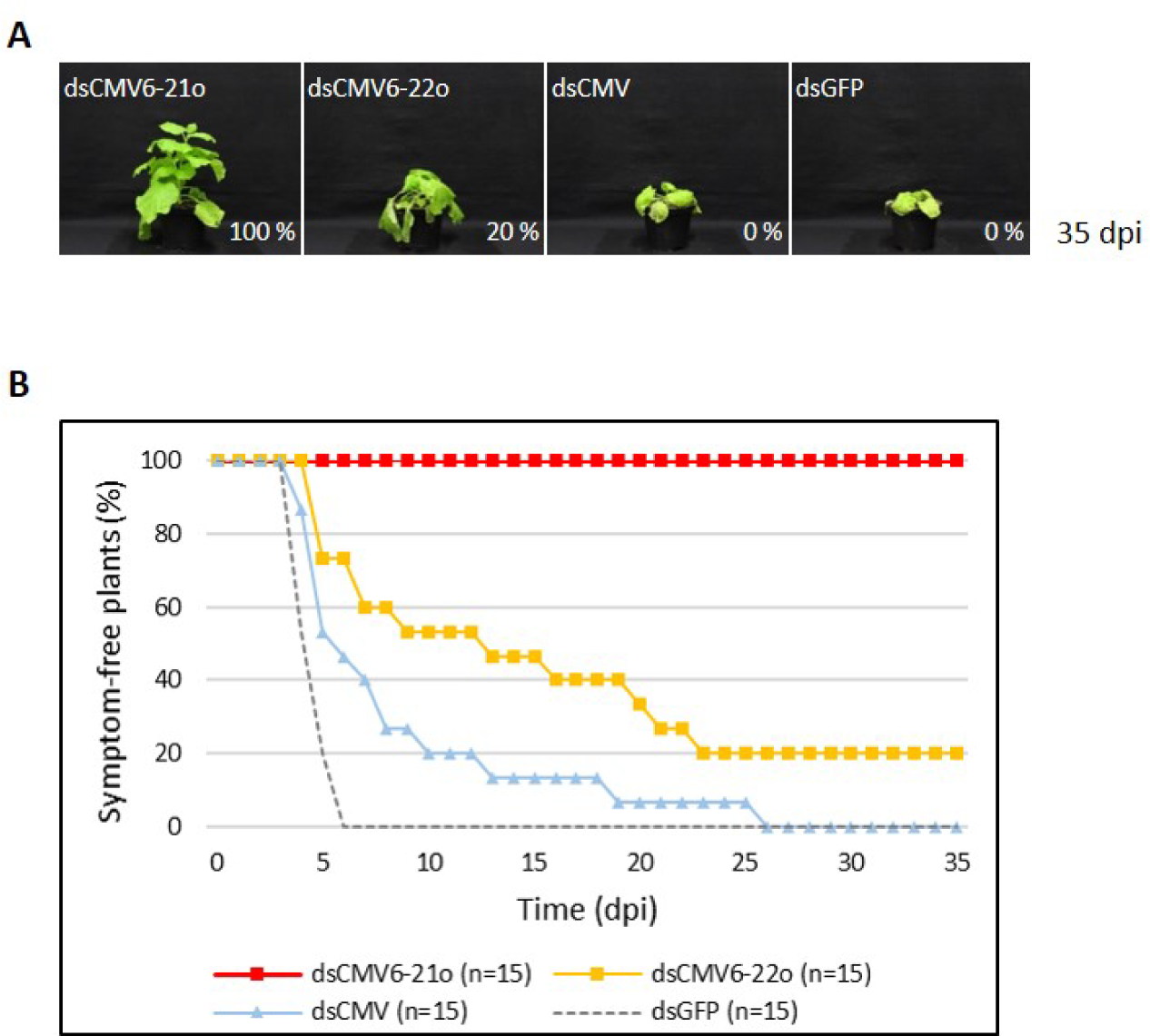
Comparison of the protective effect of different dsRNAs *in planta*. *N. benthamiana* plants were mechanically inoculated with different dsRNAs (see text) as well as the genomic CMV RNAs 1, 2, and 3 and monitored for the appearance of CMV-specific symptoms for 35 dpi. Results are from two independent experiments; the total number of plants tested is indicated (n=15). (**A**) Representative plant images 35 days after co-inoculation. The percentage of asymptomatic plants remaining is indicated for each dsRNA. **(B)** Percentage of asymptomatic plants over the entire course of the experiment.

Interestingly, treatment with dsCMV provided some protection and resulted in a delayed course of infection compared to negative controls in which plants were treated with dsGFP; however, no plants remained uninfected (without symptoms) at 35 dpi. The trend was significantly better for plants with dsCMV6-22o treatment with 20% remaining protected throughout the experiment. The most signficant results were obtained when the plants were treated with dsCMV6-21o. In this case, 80-100% of the plants remained protected against a CMV infection (**Figure 6** and **Supplementary Figure 7B**). Further control experiments clearly excluded the possibility that the observed effects were caused by contamination of the dsRNA preparations with non-hybridized single-stranded components, which could theoretically hybridize with viral RNAs during inoculation and interfere with its translation or replication (**Supplementary Figure 7B**).

Taken together, these data convincingly demonstrate the validity of our approach to develop efficient RNA actives against a devastating virus such as CMV. Our study identifies functional, highly efficacious antiviral *e*siRNAs in a first step and applies *e*dsRNAs consisting of these *e*siRNAs in a second step (**Figure 7**).

**Figure 7.**
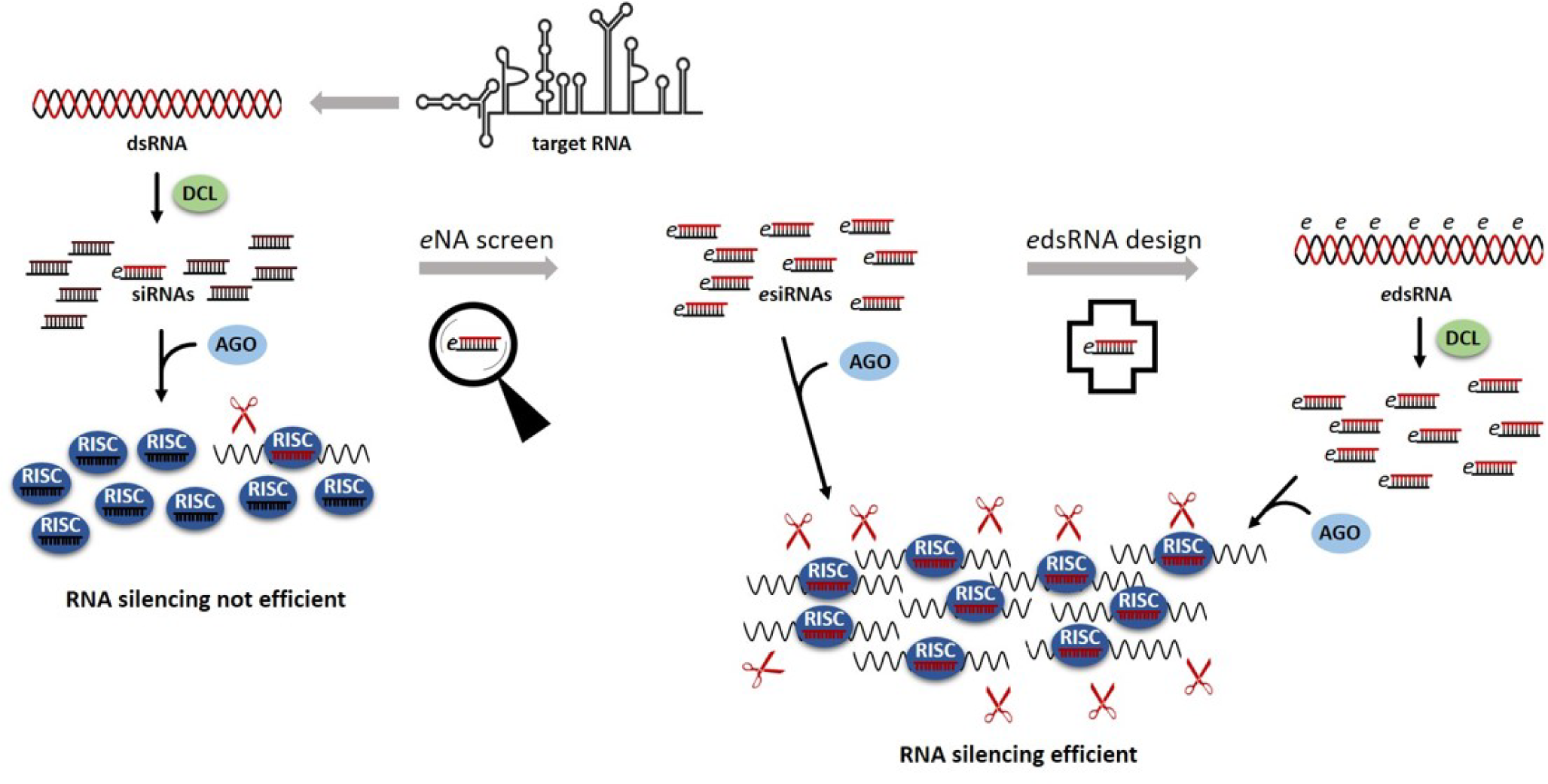
Summary of the procedures leading to the identification of *e*siRNAs and the design of *e*dsRNAs for significantly improved antiviral plant protection. Upon infection of a plant with an RNA virus, viral dsRNA (*e.g.* replication intermediates) is detected by DCLs and processed into a pool of siRNAs. Of these, only a few mediate efficient slicing of viral target RNAs by RNA-induced silencing complexes (RISC). The RNA silencing process is inefficient. Functional siRNAs, *e*siRNAs, are characterized by a high affinity to antiviral AGO proteins and high accessibility of the respective complementary target sites in the viral RNA. The *e*NA screen reliably identifies *e*siRNAs from siRNA pools. The RNA silencing process is effective; *e*siRNAs provide highly effective protection against viral infection and can be used to design *e*dsRNAs. The *e*dsRNAs are essentially composed of the sequences of the functionally characterized *e*siRNAs and are preferentially processed by the DCLs into exactly these *e*siRNAs. Multivalent *e*dsRNAs thus have the potential to significantly increase the efficiency of RNA silencing-mediated protection of plants against pathogens such as RNA viruses.

## Discussion

“RNA actives” stimulate the silencing response and are of increasing interest for the control of plant pathogens and pests that destroy up to 30% of important staple crops each year and cause enormous economic loss (73). This holds true not only for viruses, but also for fungi, nematodes and insects, where the RNA-silencing machinery can be programmed to target their mRNAs. However, considerable optimization is required to obtain maximally effective RNA actives, as a significant number of siRNAs produced by DCLs from target RNAs have no anti-pathogenic effect. Important steps toward this goal were the reconstitution of the cellular processes underlying RNA silencing *in vitro* and the application of this system to the empirical identification of *e*siRNAs that can mediate an effective antiviral immune response (32,54,55).

Until now, use of the term “effective siRNA” has been rather obscure. It evolved from *in silico* approaches that proposed RNA regions as siRNA targets because they encode conserved protein domains, or that attempted to predict RISC binding sites based on calculated RNA structures (74–76). However, these predictions are associated with large uncertainties (reviewed in (77,78)), as RNA structures are essentially functionally defined and can therefore only be verified by complex empirical studies, usually in target cells or organisms (79,80). The *in vitro* system based on cytoplasmic extracts derived from plant cells, BYL, appears to be well-suited for this purpose. Our data suggest that target RNA folding variations under these conditions are similar to those in functional plant cells. During the *e*NA screen, identified *e*siRNAs could be simultaneously functionally characterized, namely by the affinity of their guide strands to the AGO protein used (12,13) and by the successful interactions of the RISCs with a-sites of the target RNA (32-34). In other words, the *e*siRNA candidates were determined by their activity, a reproducibly quantifiable AGO1- or AGO2/RISC-mediated hydrolysis of the respective target RNA *in vitro* (**Figure 2**; **Supplementary Figure 1C**; **Tables 1** and **2**). Most importantly, the majority of these candidates actually showed a convincing protective effect against CMV when applied topically to plants, with certain *e*siRNAs reproducibly protecting 100% of individuals from infection. (**Figure 3**). It should be reiterated that the infections were carried out under an extremely strict regime. Using the mechanical “rub-inoculation” method, normally 100% of the plants developed symptoms if left untreated. The fact that such a high level of protection against CMV was achieved under these conditions, which rarely or never occur in nature in this severity, illustrates the success of the *e*NA screening procedure used to identify antiviral *e*siRNAs (**Figures 3** and **6**).

In general, a correlation was found between slicing activity *in vitro* and the antiviral efficacy of the *e*siRNA candidates *in planta*. This was most evident for the *e*siRNAs identified against CMV RNA 2 and was previously observed in the characterization of *e*siRNAs against the CMV-unrelated TBSV (32). This was less obvious for CMV RNA 3, as the number of identified *e*siRNA candidates, as well as their slicing activities and protective effects in plants were lower overall. As RNA 3 was generally inefficiently cleaved in the slicer assays, it can be assumed that the folding of this RNA generally results in poorer accessibility to RISC. The expression levels of the replicase proteins 1a and 2a are low in CMV-infected cells compared to those of the movement (3a) and capsid (3b) proteins (42). RNA 3 therefore appears to be translated at a higher rate than RNA 2, making it also more difficult for RISC to access. In addition, in CMV-infected cells, RNA 3 (and its subgenomic RNA 4) was found to be more abundant than RNA 2 (49). RNA 2 may therefore be a better target for RNA silencing, especially as the encoded RNA-dependent RNA polymerase is the earliest protagonist of viral replication (81). This idea is supported by the fact that the *e*siRNAs with the highest antiviral activity originate from the RNA 2-encoded 2a ORF. Interestingly, our *e*NA screens identified virtually no *e*siRNAs in the conserved terminal non-translated regions (NTRs) of the viral RNAs (**Figure 2; Supplementary Figure 1B**). These regions therefore appear to be particularly protected from RNA silencing, possibly due to their intense folding and functional activity in translation and replication (45,82). Taken together, our data reassert that the accessibility of functional RNA molecules to the silencing machinery can only be determined empirically.

CMV isolates are classified into three major subgroups, including IA and II, which are found worldwide, and IB, which is primarily found in East Asia (42). CMV Fny strain used in this study belongs to subgroup IA, and the *e*NA screens identified *e*siRNAs capable of targeting very different regions of RNAs 2 and 3. **Table 4** shows the sequence complementarity of the *e*siRNA candidates identified from CMV Fny to RNAs 2 or 3 of randomly selected other strains of CMV subgroups IA (O, Y, Kor, I17F), IB (Rb, Nt9) and II (Trk7, LS), respectively. As anticipated, most homologies were found in the IA strains, less in the IB strains, and least of all in the II strains. It is known that the efficacy of RISC is reduced by one or more mismatches in the seed sequence, the primary interaction region of an siRNA or microRNA with the target RNA during the AGO/RISC-mediated silencing process (40), as well as by mismatches at the central positions 9-11 located opposite the cleavage site in the target RNA (83,84). On the other hand, it has been found in the plant system that microRNAs with a few mismatches in their 3’-region to a target RNA are often as effective in RNA silencing as miRs with full complementarity, which is probably also the case for siRNAs (83). Taking all these factors into account, it is clear that most of the identified *e*siRNAs should be effective against many CMV strains of subgroups IA and IB and some also against subgroup II strains (**Table 4**). Thus, by using a combination of different *e*siRNAs targeting a-sites in different segments of the viral genome, broad-spectrum protection could be achieved, which is an important measure to counteract antigenic drifts or shifts of CMV (see also below).

**Table 4.** Protection potential of the *e*siRNA candidates identified in this study with regard to the selected CMV strains O, Y, Kor, I17F, Rb, Nt9, Trk7, and LS from different subgroups.

| Subgroup IA |  |  |  |  |  |  |  |
| --- | --- | --- | --- | --- | --- | --- | --- |
| O |  | Y |  | Kor |  | I17F |  |
| RNA 2 | RNA 3 | RNA 2 | RNA 3 | RNA 2 | RNA 3 | RNA 2 | RNA 3 |
| 149 | 35 (1) | 149 | 35 (3) | 149 (2) | 35 | 149 | 35 (1) |
| 186 | 151 | 186 | 151 | 186 | 151 | 186 | 151 |
| 359 | 239 | 359 | 239 | 359 | 239 | 359 | 239 |
| 380 | 358 | 380 | 358 | 380 | 358 | 380 | 358 |
| 407 | 478 (1) | 407 | 478 (2) | 407 | 478 (2) | 407 | 478 (1) |
| 449 | 496 | 449 | 496 (1) | 449 | 496 (1) | 449 | 496 |
| 540 | 507 (1) | 540 | 507 (1) | 540 | 507 | 540 | 507 |
| 557 | 592 | 557 | 592 | 557 | 592 (1) | 557 | 592 |
| 1020 | 593 | 1020 (1) | 593 | 1020 (2) | 593 (1) | 1020 | 593 |
| 1054 (1) | 733 | 1054 | 733 | 1054 | 733 | 1054 | 733 |
| 1172 | 985 | 1172 | 985 | 1172 | 985 | 1172 | 985 |
| 1248 | 988 | 1248 (1) | 988 | 1248 | 988 | 1248 | 988 |
| 1489 | 1019 (1) | 1489 | 1019 (1) | 1489 | 1019 | 1489 (1) | 1019 |
| 1613 (2) | 1098 | 1613 | 1098 | 1613 | 1098 (1) | 1613 | 1098 |
| 1982 | 1394 | 1982 | 1394 (1) | 1982 | 1394 | 1982 | 1394 |
| 2041 (1) | 1569 (1) | 2041 | 1569 | 2041 | 1569 (1) | 2041 | 1569 |
| 2441 |  | 2441 |  | 2441 |  | 2441 |  |
| 2562 |  | 2562 |  | 2562 |  | 2562 |  |
| 2727 |  | 2727 |  | 2727 (1) |  | 2727 |  |

| Subgroup IB |  |  |  | Subgroup II |  |  |  |
| --- | --- | --- | --- | --- | --- | --- | --- |
| Rb |  | Nt9 |  | Trk7 |  | LS |  |
| RNA 2 | RNA 3 | RNA 2 | RNA 3 | RNA 2 | RNA 3 | RNA 2 | RNA 3 |
| 149 (1) | 35 | 149 (1) | 35 | 149 (7) | 35 (m) | 149 (6) | 35 (m) |
| 186 | 151 | 186 (2) | 151 | 186 (7) | 151 (1) | 186 (7) | 151 (2) |
| 359 | 239 | 359 | 239 (1) | 359 (7) | 239 (1) | 359 (m) | 239 (1) |
| 380 | 358 | 380 (2) | 358 (1) | 380 (6) | 358 (3) | 380 (m) | 358 (3) |
| 407 | 478 (1) | 407 (1) | 478 (2) | 407 (1) | 478 (6) | 407 (1) | 478 (6) |
| 449 | 496 (1) | 449 (1) | 496 (3) | 449 (m) | 496 (4) | 449 (m) | 496 (3) |
| 540 | 507 (2) | 540 | 507 (5) | 540 (5) | 507 (4) | 540 (m) | 507 (3) |
| 557 | 592 | 557 | 592 | 557 (7) | 592 (5) | 557 (m) | 592 (5) |
| 1020 (2) | 593 | 1020 (1) | 593 | 1020 (7) | 593 (5) | 1020 (7) | 593 (5) |
| 1054 | 733 (1) | 1054 (3) | 733 (1) | 1054 (7) | 733 (4) | 1054 (7) | 733 (4) |
| 1172 | 985 (1) | 1172 (1) | 985 (3) | 1172 (4) | 985 (m) | 1172 (4) | 985 (m) |
| 1248 | 988 (1) | 1248 (1) | 988 (2) | 1248 (m) | 988 (m) | 1248 (m) | 988 (m) |
| 1489 | 1019 | 1489 (2) | 1019 | 1489 (7) | 1019 (m) | 1489 (7) | 1019 (m) |
| 1613 | 1098 | 1613 (2) | 1098 (1) | 1613 (8) | 1098 (6) | 1613 (8) | 1098 (6) |
| 1982 (1) | 1394 | 1982 | 1394 (1) | 1982 (4) | 1394 (9) | 1982 (4) | 1394 (10) |
| 2041 | 1569 | 2041 | 1569 (3) | 2041 (2) | 1569 (3) | 2041 (2) | 1569 (1) |
| 2441 |  | 2441 (3) |  | 2441 (m) |  | 2441 (m) |  |
| 2562 |  | 2562 (1) |  | 2562 (3) |  | 2562 (2) |  |
| 2727 (1) |  | 2727 (2) |  | 2727 (3) |  | 2727 (3) |  |

(...) = number of mismatches between siRNA guide strand and target sequence
(...) = at least one of the mismatches affects the siRNA's seed sequence (nt 2-8) or central positions (nt 9-11)
(m) = multiple mismatches including insertions and/or deletions
Grey background = siRNA expected to mediate cleavage of the related target RNA with similar efficiency as the cleavage of the corresponding CMV Fny RNAs.

For the optimal use of RNA actives, it is obvious that such broad-spectrum protection can best be realized in the form of applied multivalent “*e*dsRNAs” whose sequences consist largely of several *e*siRNAs and which are processed by the plant’s DCLs into precisely these *e*siRNAs. Based on studies showing that dsRNAs longer than 130 nt induce RNA silencing up to 400-fold more strongly than 21 nt- or 37 nt-long dsRNAs (85), we here established *e*dsRNA molecules with a length of ca. 170 nt. These consisted of six *e*siRNA sequences and also had other important properties **(Supplementary Figure 5**). The selection of *e*siRNAs to serve as *e*dsRNA building blocks was based on their functionality, *i.e.,* their preferential incorporation into AGO1- or AGO2/RISC and their antiviral efficacy (**Table 1**). According to a model by Harvey et al. (86), the RNA-silencing defense of the plant against an infecting virus has several levels, with AGO1 being part of a first layer and AGO2 part of a second. In uninfected plants, the expression of AGO1 is usually higher than that of AGO2 (87), and there is evidence that miR403-loaded AGO1/RISC downregulate AGO2 mRNA expression post-transcriptionally (88). In virus-infected *N. benthamiana*, an increase in AGO2 levels can be detected (89), and one model to explain this is that viral suppressors of RNA silencing, VSRs, sequester miR403 and thus override the suppression of AGO2 expression by AGO1 (57). Consistent with this, VSRs such as the CMV 2b have been shown to bind small RNAs with high affinity and also directly inhibit AGO1 activity (90). Thus, the activation of the second layer of the plant cell’s RNA-silencing defense appears to be a direct consequence of the loss of the first layer (86); when AGO1 activity is blocked, AGO2 is an important backup to limit viral accumulation. The construction of *e*dsRNAs from AGO1- and AGO2-incorporating *e*siRNAs should therefore address both levels of the plant’s antiviral RNAi immune response to achieve maximum antiviral efficacy.

The second important feature of the “*e*dsRNA active” toolkit established in this study concerned the “pseudo-siRNA sequences” at the termini (**Figure 4A** and **Supplementary Figure 5**). On the DNA-level these sequences enabled high yield transcription. On the RNA-level, they were intended to support a phased processing by the DCLs by providing the appropriate distance between the 5’-ends of the dsRNA and the 5’-terminal siRNA sequence. Our data show that the pseudo-siRNA sequences do indeed serve both functions. Above all, we were able to confirm that the *e*dsRNAs are apparently predominantly processed by the DCLs in such a way that large quantities of the individual *e*siRNAs are actually produced. For example, an *e*dsRNA with 21 nt pseudo-siRNA-termini generates large quantities of 21 nt *e*siRNAs, as predicted (**Figure 5B**). The success of this approach was demonstrated in the plant-protection experiments, whereby, compared to a classical dsRNA (dsCMV), which happened to contain two *e*siRNA sequences, the antiviral effect of the multivalent *e*dsRNAs was significantly higher (**Figure 6**). While the conventional dsRNA only delayed the development of symptoms, and thus the infection process without providing protection, the *e*dsRNAs protected 80-100% of the infected plants.

Previous reports have shown that 21 nt siRNAs predominate over 22 nt siRNAs in CMV-infected plants (16,91-94) and, as in other plant virus infection systems, evidence for a redundant role of DCL4 and DCL2 has been presented (95,96). Further observations indicated that efficient secondary siRNA-dependent silencing of CMV-Δ2b in *A. thaliana* relies on DCL4-dependent 21 nt siRNAs, but not on DCL2-dependent 22 nt siRNAs. The latter were insufficient to confer resistance to the virus in the absence of DCL4 (16). Overall, this suggests that the processing of dsRNAs by DCL4 and the generation of 21 nt siRNAs are particularly important for the induction of an antiviral RNA-silencing response against CMV. Our *in vitro* and *in planta* data support this proposal. They show that 21 nt-long *e*siRNAs and *e*dsRNAs containing 21 nt *e*siRNA sequences are more efficient in the antiviral RNA-silencing process than 22 nt *e*siRNAs and *e*dsRNAs containing 22 nt *e*siRNA sequences (**Figure 6** and **Supplementary Figure 4**). We have no evidence to indicate that the 22 nt *e*siRNAs used in this study enhanced CMV silencing by inducing secondary siRNA production.

The third feature of the “*e*dsRNA actives” developed here concerned optional 3’-overhangs. Interestingly, we observed no differences in antiviral activity when we used blunt or 3’ overhanging *e*dsRNAs in the *in vitro* and infectious virus-plant systems, respectively. This suggests that the DCLs involved in the processing of *e*dsRNAs to *e*siRNAs, namely DCL4 and/or DCL2, do not have substrate preferences that depend on the presence of 3’-overhangs. However, it is important to note here that when *e*dsRNAs are considered for use in other organisms, the situation may be different. For example, the activity, including the phased processing, of Dicer on dsRNAs in insects appears to be largely affected by the nature of the dsRNA termini (72).

The processing pattern of *e*dsRNAs in BYL clearly showed that in addition to the expected *e*siRNAs, siRNAs of other lengths, particularly those with a length of 24 nt, were generated (**Figure 5A**). The presence of other RNases in these extracts, above all DCL3, may explain this, but imperfectly phased processing by DCL4 and/or DCL2 could also be a contributing factor. Although this was not tested, it is expected that the use of *e*dsRNAs may not prevent the production of unwanted siRNAs in plants. However, it is clear that, compared to classically organized dsRNAs, the off-target effects of *e*dsRNA-generated siRNAs on mRNAs of the host plant (31,38), as well as on non-target organisms (41,97), should be significantly reduced. One of the reasons for this is the large number and quantity of *e*siRNAs that are generated, as mentioned above. On the other hand, according to our data and the underlying principles of *e*NA screening, it can be assumed that *e*siRNAs are incorporated into RISC at considerably higher rates than other siRNAs generated from *e*dsRNAs due to their high affinity for AGO1 or AGO2. In addition, we assume that significantly lower amounts of *e*dsRNA are required for topical application to achieve a protective effect compared to conventional dsRNA. Accordingly, using *e*dsRNA actives in topical RNA-silencing approaches will reduce the risk of decoy and off-target effects. The results of this study therefore offer the potential for the efficient use of RNA in the biological protection of plants. It will now be important to further test and further improve upon the topical RNA applications achieved in this study in combination with suitable dsRNA formulations in agricultural applications, such as in greenhouse or field experiments. Our data already suggest that multivalent *e*dsRNAs are capable to induce RNA silencing at multiple RNA targets. This suggests that in the future, single “*e*dsRNA actives” may be developed that target multiple pathogens simultaneously.

## Data availability

NGS datasets have been deposited in the ENA database under accession number PRJEB76219.

## Supplementary data

Supplementary data are available at online.

## Acknowledgement

We thank Gary Sawers for critical reading of the manuscript and Christine Hamann and Katja Rostowski for technical support. We thank the Leibniz Institute of Plant Biochemistry (IPB) in Halle (Saale) and its gardeners for providing *N. benthamiana* plants. We are grateful to Prof. Fernando García-Arenal Rodríguez (Universidad Politécnica de Madrid) and Prof. John Carr (University of Cambridge) for providing the CMV Fny full-length cDNA clones.

