## Supplementary data for "A new level of RNA-based plant protection - dsRNAs designed from functionally characterized siRNAs highly effective against Cucumber Mosaic Virus"

**Supplementary Materials and methods**

**Analysis of DCL-mediated processing of *e*dsRNA**

200 ng ^32^P-labeled RNA was incubated for up to 24 h at 25°C in 20 µl reactions containing 50% (v/v) BYL and the buffer conditions described above. Total RNA was isolated from the reaction by treatment with proteinase K (Thermo Scientific, #EO0491), followed by chloroform extraction and ethanol precipitation. RNAs were separated on 15% denaturing urea-polyacrylamide gels in TBE buffer and siRNAs were visualized by phosphor-imaging (Typhoon Trio+, GE Healthcare, Chalfont St Giles, UK). To analyze the siRNAs resulting from processing of the *e*dsRNA by RNA-Seq, 1.25 µg RNA was incubated for 4 h in a 50 µl reaction containing 50% (v/v) BYL using the aforementioned conditions. Isolation of total RNA and RNA-Seq was performed as described above.

**Western Blotting**

For the immunodetection of *in vitro* synthesized, FLAG-tagged AGO proteins, samples were separated on 10% polyacrylamide gels and western blotting was performed at standard conditions using semi-dry transfer. Polyclonal anti-FLAG-tag antibody (Sigma-Aldrich, #F7425-.2MG) was used at a 1:1000 dilution and detected by IRDye 800CW goat anti-rabbit IgG secondary antibody (LI-COR, Lincoln, NE, #926-32211) using a LI-COR ODYSSEY CLx imager.

**Silver staining of RNAs**

siRNAs that were co-purified in AGO immunoprecipitations were separated on 15% (w/v) denaturing urea-polyacrylamide gels and visualized by silver staining according to Blum et al. (1987). The gel was first fixed in 50% (v/v) methanol, 12% (v/v) acetic acid, and 0.0185% (v/v) formaldehyde for 1 h and subsequently washed three times for 20 min each in 50% (v/v) ethanol. Following pre-treatment with 0.02% (w/v) Na_2_S_2_O_3_ × 5 H_2_O for 1 min, the gel was rinsed three times for 20 s with water and incubated for 20 min in a solution containing 0.2% (w/v) AgNO_3_ and 0.028% (v/v) formaldehyde. After rinsing twice with water for 20 s each time, the image development was performed in 6% (w/v) Na_2_CO_3_, 0.0185% (v/v) formaldehyde, and 0.0004% (w/v) Na_2_S_2_O_3_ × 5 H_2_O for 10 min. The gel was washed twice with water for 2 min each time and the staining reaction was stopped by incubating for 10 min in a solution of 50% (v/v) methanol and 12% (v/v) acetic acid. Finally, the gel was washed in 50% (v/v) MeOH for at least 20 min.

**Monitoring of viral RNA replication**

RNA isolated from CMV-inoculated plants was treated with RNase-free DNase I (Roche Diagnostics, #04716728001) and reverse-transcribed using random hexamer primers and RevertAid M-MuLV Reverse Transcriptase (Thermo Scientific, #EP0441) according to the manufacturer's instructions. PCR amplification was performed using DreamTaq DNA Polymerase (Thermo Scientific, #EP0705) and a primer pair that binds in the 3’ region of each of the three CMV RNAs.

**Supplementary Tables**

**Supplementary Table 1.** List of oligonucleotides

| **DNA oligonucleotides** | | |
| --- | --- | --- |
| Name | Sequence (5'-3') | Purpose |
| T7CMV2_MK1f | CCCTAATACGACTCACTATAGGTTTATTTACAAGAGCGTAC | PCR primer for the generation of template DNA for *in vitro* transcription of CMV RNA 1 and 2, (+)-strand |
| TraCMV1T_MK1r | TGGTCTCCTTTTAGAGACC | PCR primer for the generation of template DNA for *in vitro* transcription of CMV RNA 1, (+)-strand |
| TraCMV2T_MK1r | TGGTCTCCTTTTGGAGGCC | PCR primer for the generation of template DNA for *in vitro* transcription of CMV RNA 2 and 3, (+)-strand |
| T7CMV3_MK1f | CCCTAATACGACTCACTATAGGTAATCTTACCACTGTGTGTG | PCR primer for the generation of template DNA for *in vitro* transcription of CMV RNA 3, (+)-strand |
| TraCMV2G_MK1r | GGGTCTCCTTTTGGAGGCC | PCR primer for the generation of template DNA for *in vitro* transcription of CMV RNA 2 and 3, (+)-strand with 3' terminal C |
| T7CMV2_MK1r | CCCTAATACGACTCACTATAGGGTCTCCTTTTGGAGGCC | PCR primer for the generation of template DNA for *in vitro* transcription of CMV RNA 2 and 3, (-)-strand with 5’ terminal G |
| TraCMV2_MK1f | GGTTTATTTACAAGAGCGTAC | PCR primer for the generation of template DNA for *in vitro* transcription of CMV RNA 2, (-)-strand |
| TraCMV3_MK1f | GGTAATCTTACCACTGTGTGTG | PCR primer for the generation of template DNA for *in vitro* transcription of CMV RNA 3, (-)-strand |
| T7VSVdsTra1f | GTCTTTCAGGAAAAAAACTAACAGATATCATGGATATGCTAGGTAATACGACTCACTATAGGGGGTCTTC | oligonucleotide for generating a modified pUC18 vector that contains opposite T7 promoters flanking two BpiI (BbsI) sites |
| T7VSVdsTra1r | TCGAGAAGACCCCCTATAGTGAGTCGTATTACCTAGCATATCCATGATATCTGTTAGTTTTTTTCCTGAAAGACTGCA | oligonucleotide for generating a modified pUC18 vector that contains opposite T7 promoters flanking two BpiI (BbsI) sites |
| T7VSVdsTra2f | CTAGGAAGACCCCTATAGTGAGTCGTATTACCTAGCATATCCATGATATCTGTTAGTTTTTTTCCTGAAAGAG | oligonucleotide for generating a modified pUC18 vector that contains opposite T7 promoters flanking two BpiI (BbsI) sites |
| T7VSVdsTra2r | GATCCTCTTTCAGGAAAAAAACTAACAGATATCATGGATATGCTAGGTAATACGACTCACTATAGGGGTCTTC | oligonucleotide for generating a modified pUC18 vector that contains opposite T7 promoters flanking two BpiI (BbsI) sites |
| T7VSVadapt1f | TCGAAAGCGGCCGCCCATGGTT | oligonucleotide for generating a modified pUC18 vector that contains opposite T7 promoters flanking two BpiI (BbsI) sites |
| T7VSVadapt1r | CTAGAACCATGGGCGGCCGCTT | oligonucleotide for generating a modified pUC18 vector that contains opposite T7 promoters flanking two BpiI (BbsI) sites |
| fragGFP168-1f | GACCATGAAGACTCTAGGGCGAATTGGGTACCGGGCAAGATCTGAGTCCGGACTTGT | PCR primer for cloning of a GFP gene fragment used as template for the generation of an unspecific control dsRNA |
| fragGFP168-1r | GACCATGAAGACTCATAGGGAGACCGGCAGATCTGATGTCCACACAATCTGCCCTTTC | PCR primer for cloning of a GFP gene fragment used as template for the generation of an unspecific control dsRNA |
| fragCMV168-1f | GACCATGAAGACTCTAGGGCGAATTGGGTACCGGGGAATAACCGGGTACATCGCGAG | PCR primer for cloning of a CMV RNA 2 cDNA fragment used as template for the generation of a control dsRNA |
| fragCMV168-1r | GACCATGAAGACTCATAGGGAGACCGGCAGATCTGAACCGAACCATGAAGTGTTTT | PCR primer for cloning of a CMV RNA 2 cDNA fragment used as template for the generation of a control dsRNA |
| siR6CMV21-1f | TAGGGCGAATTGGGTACCGGGCATTACGTTTCTTAATTGCTGTATACTCTTCTTATGATACGTTATCAGATTTTTCAAGGTAATCT | oligonucleotide for cloning of CMV RNA 2 cDNA fragments used as template for the generation of an edsRNA consisting of six 21 nt esiRNAs |
| siR6CMV21-2f | TGATGAGCTCCTTGTCGCTTTTTTGTTCGAAGTTCTTACTCTTTCGTCGAAAGTGCAGACTATTCTCAGATCTGCCGGTCTCC | oligonucleotide for cloning of CMV RNA 2 cDNA fragments used as template for the generation of an edsRNA consisting of six 21 nt esiRNAs |
| siR6CMV21-1r | ATCAAGATTACCTTGAAAAATCTGATAACGTATCATAAGAAGAGTATACAGCAATTAAGAAACGTAATGCCCGGTACCCAATTCGC | oligonucleotide for cloning of CMV RNA 2 cDNA fragments used as template for the generation of an edsRNA consisting of six 21 nt esiRNAs |
| siR6CMV21-2r | ATAGGGAGACCGGCAGATCTGAGAATAGTCTGCACTTTCGACGAAAGAGTAAGAACTTCGAACAAAAAAGCGACAAGGAGCTC | oligonucleotide for cloning of CMV RNA 2 cDNA fragments used as template for the generation of an edsRNA consisting of six 21 nt esiRNAs |
| siR6CMV22-1f | TAGGGCGAATTGGGTACCGGGCCATTACGTTTCTTAATTGCTGTAATACTCTTCTTATGATACGTTATTCAGATTTTTCAAGGTAATCTT | oligonucleotide for cloning of CMV RNA 2 cDNA fragments used as template for the generation of an edsRNA consisting of six 22 nt esiRNAs |
| siR6CMV22-2f | ATGATGAGCTCCTTGTCGCTTTATTTGTTCGAAGTTCTTACTCTGTTCGTCGAAAGTGCAGACTATTCATCAGATCTGCCGGTCTCC | oligonucleotide for cloning of CMV RNA 2 cDNA fragments used as template for the generation of an edsRNA consisting of six 22 nt esiRNAs |
| siR6CMV22-1r | TCATAAGATTACCTTGAAAAATCTGAATAACGTATCATAAGAAGAGTATTACAGCAATTAAGAAACGTAATGGCCCGGTACCCAATTCGC | oligonucleotide for cloning of CMV RNA 2 cDNA fragments used as template for the generation of an edsRNA consisting of six 22 nt esiRNAs |
| siR6CMV22-2r | ATAGGGAGACCGGCAGATCTGATGAATAGTCTGCACTTTCGACGAACAGAGTAAGAACTTCGAACAAATAAAGCGACAAGGAGCTCA | oligonucleotide for cloning of CMV RNA 2 cDNA fragments used as template for the generation of an edsRNA consisting of six 22 nt esiRNAs |
| SeqUPN | AGGGTTTTCCCAGTCACGACGTTG | vector-specific PCR primer for the generation of template DNA for *in vitro* transcription of the top strand of edsRNAs and control dsRNAs |
| TrasiR6CMV-A1-1r | GGGAGACCGGCAGATCTG | PCR primer for the generation of template DNA for *in vitro* transcription of the top strand of edsRNAs and control dsRNAs with blunt ends |
| TrasiR6CMV-A1-2r | TAGGGAGACCGGCAGATCTG | PCR primer for the generation of template DNA for *in vitro* transcription of the top strand of edsRNAs and control dsRNAs with 2 nt 3’ overhang |
| SeqRPN | CAATTTCACACAGGAAACAGCTATG | vector-specific PCR primer for the generation of template DNA for *in vitro* transcription of the bottom strand of edsRNAs and control dsRNAs |
| TrasiR6CMV-A2-1r | GGGCGAATTGGGTACCGG | PCR primer for the generation of template DNA for *in vitro* transcription of the bottom strand of edsRNAs and control dsRNAs with blunt ends |
| TrasiR6CMV-A2-2r | TAGGGCGAATTGGGTACCGG | PCR primer for the generation of template DNA for *in vitro* transcription of the bottom strand of edsRNAs and control dsRNAs with 2 nt 3’ overhang |
| T7_CMV-S_3'reg1r | CCCTAATACGACTCACTATAGGGCACCCGTACCCTGAAACTAGC | PCR primer for the detection of CMV infection by RT-PCR |
| Tra_CMV-S_3'reg1f | GGCGGGATCTGAGTTGGC | PCR primer for the detection of CMV infection by RT-PCR |
| **RNA oligonucleotides^a^** | | |
| Name | Sequence (5'-3') | Purpose |
| siR gf698-21U gs | uaguucauccaugccaugugu | guide strand of siRNA gf698, 5' U |
| siR gf698-21U ps | acauggcauggaugaacuaua | passenger strand of siRNA gf698, (5' U in gs) |
| siR gf698-21A gs | aaguucauccaugccaugugu | guide strand of siRNA gf698, 5' A |
| siR gf698-21A ps | acauggcauggaugaacuuua | passenger strand of siRNA gf698, (5' A in gs) |
| siR359-22 gs | ucagauuuuucaagguaaucuu | guide strand of 22 nt variant of siRNA 359 |
| siR359-22 ps | gauuaccuugaaaaaucugaug | passenger strand of 22 nt variant of siRNA 359 |
| siR380-22 gs | aaagcgacaaggagcucaucau | guide strand of 22 nt variant of siRNA 380 |
| siR380-22 ps | gaugagcuccuugucgcuuuug | passenger strand of 22 nt variant of siRNA 380 |
| siR1020-22 gs | auagucugcacuuucgacgaac | guide strand of 22 nt variant of siRNA 1020 |
| siR1020-22 gs | ucgucgaaagugcagacuauuc | passenger strand of 22 nt variant of siRNA 1020 |
| siR1172-22 gs | uuacguuucuuaauugcuguaa | guide strand of 22 nt variant of siRNA 1172 |
| siR1172-22 ps | acagcaauuaagaaacguaaug | passenger strand of 22 nt variant of siRNA 1172 |
| siR1489-22 gs | uacucuucuuaugauacguuau | guide strand of 22 nt variant of siRNA 1489 |
| siR1489-22 ps | aacguaucauaagaagaguaua | passenger strand of 22 nt variant of siRNA 1489 |
| siR2041-22 gs | agaguaagaacuucgaacaaau | guide strand of 22 nt variant of siRNA 2041 |
| siR2041-22 ps | uuguucgaaguucuuacucucu | passenger strand of 22 nt variant of siRNA 2041 |

^a^Sequences of RNA oligonucleotides that correspond to the 21 nt *e*siRNA candidates targeting CMV RNAs 2 and 3 are listed in tables 1 and 2, respectively

**Supplementary Table 2.** Accumulation of CMV siRNAs in AGO/RISC

| **CMV RNA 2** | | | | | |
| --- | --- | --- | --- | --- | --- |
| **AGO1 IP/DCL processed pool^a^** | | | **AGO2 IP/DCL processed pool^a^** | | |
| **siRNA** | **5’ nt** | **log2Fold change** | **siRNA** | **5’ nt** | **log2Fold change** |
| 2441 | U | 6.6 | 2634 | A | 5.6 |
| 2562 | U | 5.5 | 1054 | A | 5.3 |
| 186 | U | 5.4 | 2041 | A | 5.0 |
| 1613 | U | 4.9 | 1020 | A | 4.8 |
| 1844 | G | 4.7 | 557 | A | 4.7 |
| 1489 | U | 4.6 | 2801 | A | 4.7 |
| 149 | U | 4.5 | 2863 | A | 4.6 |
| 1982 | U | 4.5 | 2955 | A | 4.5 |
| 2727 | U | 4.5 | 1248 | A | 4.5 |
| 1172 | U | 4.5 | 380 | A | 4.5 |
| **AGO1 IP (not detected in DCL processed pool)^b^** | | | **AGO2 IP (not detected in DCL processed pool)^b^** | | |
| **siRNA** | **5’ nt** | **mean abundance (%)** | **siRNA** | **5’ nt** | **mean abundance (%)** |
| 359 | U | 0.00608 | 540 | A | 0.01022 |
| 2740 | U | 0.00608 | 449 | A | 0.00461 |
|  |  |  | 2748 | A | 0.00403 |
|  |  |  | 407 | A | 0.00389 |
| **CMV RNA 3** | | | | | |
| **AGO1 IP/DCL processed pool^a^** | | | **AGO2 IP/DCL processed pool^a^** | | |
| **siRNA** | **5’ nt** | **log2Fold change** | **siRNA** | **5’ nt** | **log2Fold change** |
| 196 | A | 6.6 | 667 | C | 7.4 |
| 507 | U | 5.0 | 496 | A | 6.4 |
| 34 | C | 4.7 | 592 | A | 5.3 |
| 33 | A | 4.7 | 593 | A | 5.3 |
| 985 | U | 4.4 | 478 | A | 5.2 |
| 239 | U | 4.1 | 1569 | A | 4.9 |
| 2061 | U | 3.8 | 1019 | A | 4.9 |
| 151 | U | 3.7 | 733 | A | 4.7 |
| 2174 | U | 3.7 | 35 | A | 4.5 |
| 1098 | U | 3.7 | 2099 | A | 4.3 |
| **AGO1 IP (not detected in DCL processed pool)^b^** | | | **AGO2 IP (not detected in DCL processed pool)^b^** | | |
| **siRNA** | **5’ nt** | **mean abundance (%)** | **siRNA** | **5’ nt** | **mean abundance (%)** |
| 988 | U | 0.00642 | 1394 | A | 0.00574 |
|  |  |  | 1132 | A | 0.00463 |
|  |  |  | 358 | A | 0.00390 |

^a^siRNAs were sorted according to the ratio (log2Fold change) of their abundance in the AGO immunoprecipitation to their abundance in the initial siRNA pool generated from dsRNAs by BYL-endogenous DCLs

^b^siRNAs listed here were not detectable in the initial siRNA pool (i.e. a log2Fold change could not be calculated) but showed in the AGO immunoprecipitation a higher abundance than at least one of the siRNAs listed above

**Supplementary Figure 1**

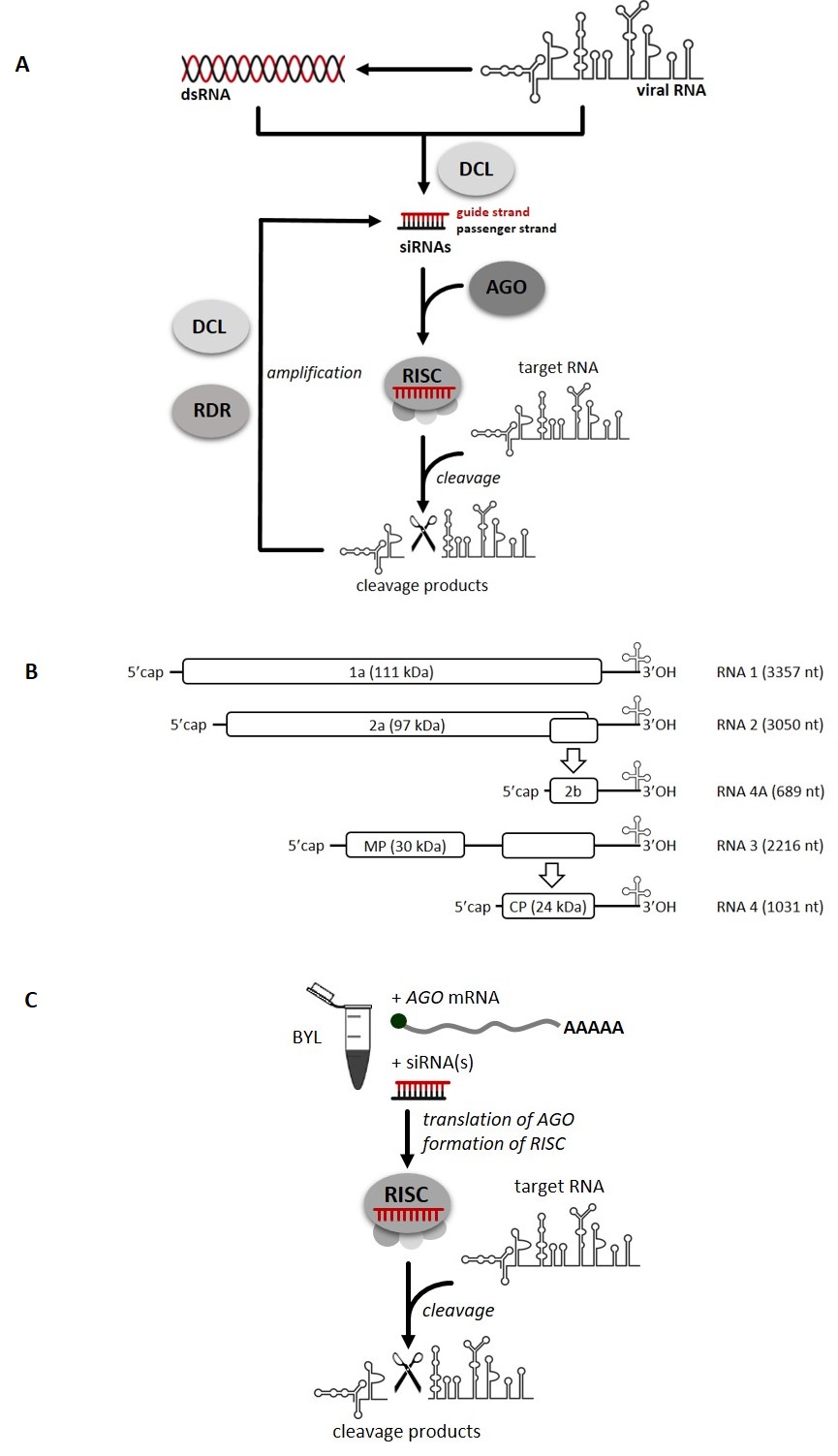

**Supplementary Figure 1. Schematic representations of the antiviral RNAi pathway in plants, the genomic organization of CMV and the *in vitro* slicer assay. (A)** Antiviral RNAi pathway that leads to the slicing (cleavage) of a viral target RNA (for a detailed description, see text). Viral genomic RNA or double-stranded (ds) versions of the viral RNA such as replication intermediates are processed by DCLs into siRNAs. SiRNA guide strands are incorporated into AGO/RISC, which then may catalyze endonucleolytic cleavage of the complementary target viral RNA. Secondary siRNAs can be generated in an amplification mechanism mediated by the activity of RDRs and DCLs (see text). **(B)** Genomic organization of the CMV genome. All viral RNAs are capped and contain a tRNA-like structure at their 3’-ends. Untranslated regions (UTRs) are shown as lines, ORFs as boxes. The names and molecular weights of the proteins encoded by the genomic (RNAs 1, 2 and 3) and subgenomic RNAs (RNAs 4A and 4) are indicated. MP, movement protein; CP, capsid protein. **(C)** *In vitro* “slicer assay” with BYL that reproduces RISC-mediated slicing (cleavage) of a target RNA with a translated AGO protein of choice. The technical details of the assay are described in the text. It is important to note that the assay can be performed with siRNAs generated endogenously from a dsRNA by the DCLs present in BYL as well as with synthetic siRNAs added exogenously.

**Supplementary Figure 2**

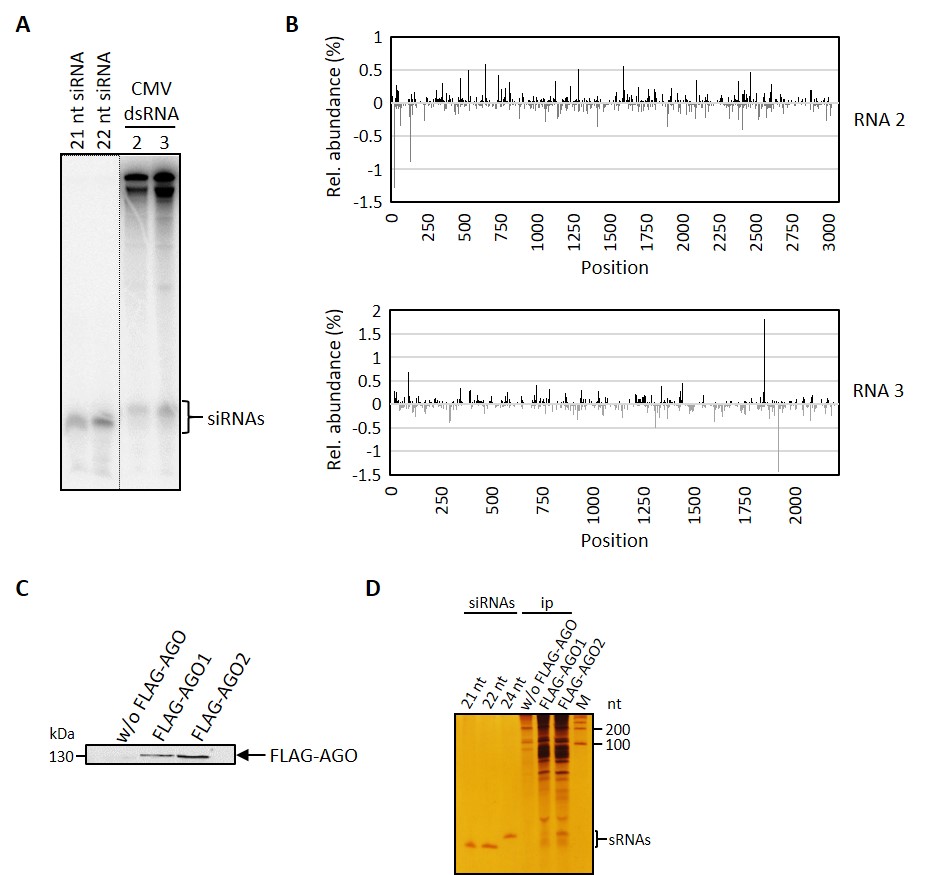

**Supplementary Figure 2. DCL-mediated processing of double-stranded versions of CMV RNAs 2 and 3 in BYL and isolation of AGO-bound siRNAs (*e*NA screen steps 1 and 2). (A)** Denaturing PAGE and autoradiography of ^32^P-labeled, double-stranded (ds) versions of CMV RNAs 2 and 3 that were processed in BYL by endogenous DCLs. Synthetic, single-stranded RNAs of a length of 21 and 22 nt served as markers. **(B)** Distribution and abundance of 21 nt siRNAs that were processed by BYL-endogenous DCLs from ds versions of CMV RNAs 2 and 3: total RNA was isolated and the small RNA fraction analyzed by RNA-seq. Peaks above and below the axis represent siRNAs derived from viral (+) strand RNA and viral (-) strand RNA, respectively. Peaks correspond to the position of the 5’-nucleotide of the respective siRNA strands. Data represent the mean values from three experiments. **(C)** Exemplary western blot of FLAG-AGO samples collected after stringent washing of the Anti-FLAG M2 affinity gel during the course of the immunoprecipitation procedure (see Supplementary Materials and methods). The precipitated FLAG-AGO proteins are indicated. **(D)** Detection of AGO-bound small RNAs. RNAs were isolated from immunoprecipitated FLAG-AGO proteins, separated by denaturing PAGE, and visualized by silver staining. Synthetic, single-stranded siRNAs of 21 nt, 22 nt, and 24 nt, respectively, were used for comparison. M = RNA molecular weight marker.

**Supplementary Figure 3**

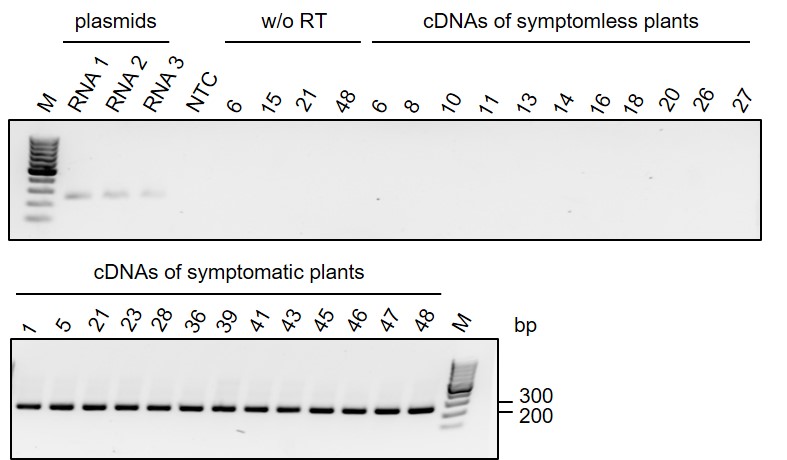

**Supplementary Figure 3. Detection of CMV infection by RT-PCR.** Leaf discs of asymptomatic and symptomatic *N. benthamiana* plants treated with siRNAs and genomic CMV RNAs (see text) were collected at 35 dpi. The numbers correspond to the numbering of the respective plants from the infection experiment performed with siRNAs directed against CMV RNA 2 (**Figure 3**). Total RNA was extracted and cDNA synthesized using Reverse Transcriptase (RT). This was followed by PCR to detect a conserved sequence in CMV RNAs 1, 2 and 3 and separation of the PCR products on an agarose gel. Amplification of the CMV-specific sequence from plasmids containing the cDNA sequence of the respective viral RNA served as positive control. Samples without addition of RT to the cDNA synthesis reaction (w/o RT) and a “no template control” (NTC, addition of water instead of cDNA during the PCR reaction) served as negative controls (M = GeneRuler 100 bp DNA ladder).

**Supplementary Figure 4**

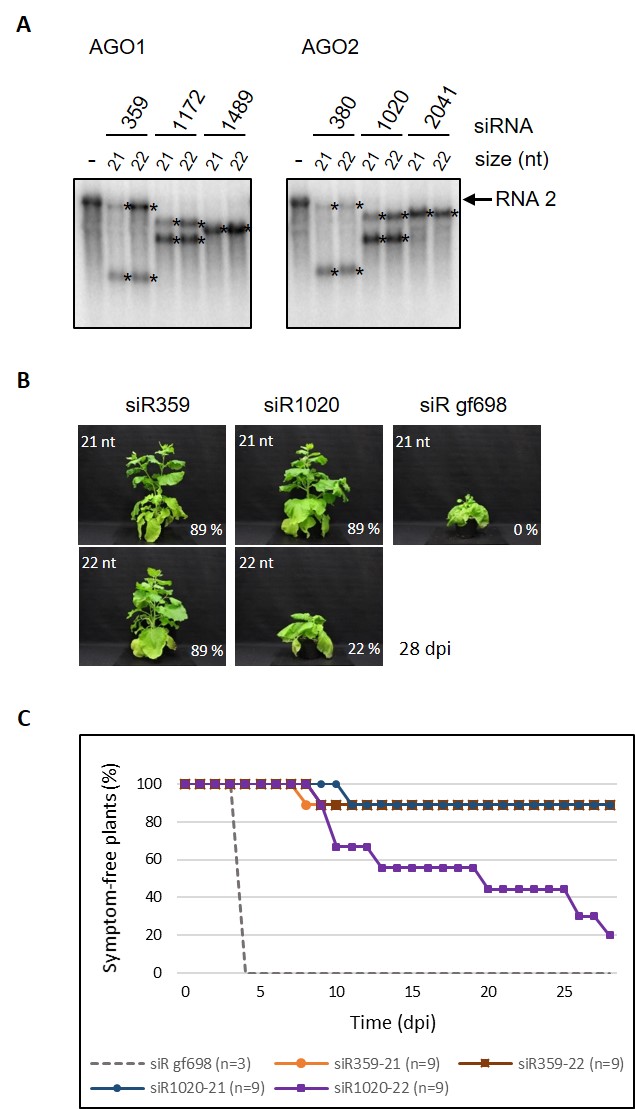

**Supplementary Figure 4. Efficacy of 21 nt and 22 nt long *e*siRNAs against CMV RNA 2 *in vitro* and *in planta*. (A)** Slicer assay using examples of 21 nt *e*siRNA as well as corresponding 22 nt variants (guide strands extended at the 3’-end). **(B)** Comparison of the protective effect of 21 nt and 22 nt siRNA *in planta*. *N. benthamiana* plants were mechanically co-inoculated with the synthetic *e*siRNAs and with the genomic CMV RNAs (see text). Representative plant images 28 days after co-inoculation are shown. The percentage of asymptomatic remaining plants is indicated for each siRNA. **(C)** Percentage of asymptomatic plants over the entire course of the experiment. The siRNA gf698 targeting GFP mRNA served as negative control. Results from one experiment including 9 plants for each CMV-targeting siRNA and 3 plants for the siR gf698 control.

**Supplementary Figure 5**

**dsCMV6-21o**

**5’GGGCGAAUUGGGUACCGGGCA**UUACGUUUCUUAAUUGCUGUAUACUCUUCUUAUGAUACGUUAUCAGAUUUUUCAAGGUAAUCUUGAUGAGCUCCUUGUCGCUUUUUUGUUCGAAGUUCUUACUCUUUCGUCGAAAGUGCAGACUAU**UCUCAGAUCUGCCGGUCUCCC**UA3**’**

3’AU**CCCGCUUAACCCAUGGCCCGU**AAUGCAAAGAAUUAACGACAUAUGAGAAGAAUACUAUGCAAUAGUCUAAAAAGUUCCAUUAGAACUACUCGAGGAACAGCGAAAAAACAAGCUUCAAGAAUGAGAAAGCAGCUUUCACGUCUGAUA**AGAGUCUAGACGGCCAGAGGG5’**

**dsCMV6-22o**

**5’GGGCGAAUUGGGUACCGGGCCA**UUACGUUUCUUAAUUGCUGUAAUACUCUUCUUAUGAUACGUUAUUCAGAUUUUUCAAGGUAAUCUUAUGAUGAGCUCCUUGUCGCUUUAUUUGUUCGAAGUUCUUACUCUGUUCGUCGAAAGUGCAGACUAU**UCAUCAGAUCUGCCGGUCUCCC**UA3’

3’AU**CCCGCUUAACCCAUGGCCCGGU**AAUGCAAAGAAUUAACGACAUUAUGAGAAGAAUACUAUGCAAUAAGUCUAAAAAGUUCCAUUAGAAUACUACUCGAGGAACAGCGAAAUAAACAAGCUUCAAGAAUGAGACAAGCAGCUUUCACGUCUGAUA**AGUAGUCUAGACGGCCAGAGGG5’**

**dsCMV**

**5’GGGCGAAUUGGGUACCGGGGA**AUAACCGGGUACAUCGCGAGACGAGAUAUCUAUCUGAGCGUCGUCGGCUUCACACUCUUCACUGUAGUAGAAAUCAGAUUUAGUGUA*AAUAGCCGCGACCAGGUCUUC*AAAACACUUCAUGGUUCG**GUUCAGAUCUGCCGGUCUCCCUA3’**

3’AU**CCCGCUUAACCCAUGGCCCCU**UAUUGGCCCAUGUAGCGCUCUGCUCUAUAGAUAGACUCGCAGCAGCCGAAGUGUGAGAAGUGACAUCAUCUUUAGUCUAAAUCACAUUUAUCGGCGCUGGUCCAGAAGUUUUGUGAAGUACCAAGC**CAAGUCUAGACGGCCAGAGGG5’**

**dsGFP**

**5’GGGCGAAUUGGGUACCGGGCA**AGAUCUGAGUCCGGACUUGUAuaguucauccaugccauguguAAUCCCAGCAGCUGUUACAAACUCAAGAAGGACCAUGUGGUCUCUCUUUUCGUUGGGAUCUUUCGAAAGGGCAGAUUGUGUGGA**CAUCAGAUCUGCCGGUCUCCC**UA3’

3’AU**CCCGCUUAACCCAUGGCCCGU**UCUAGACUCAGGCCUGAACAUAUCAAGUAGGUACGGUACACAUUAGGGUCGUCGACAAUGUUUGAGUUCUUCCUGGUACACCAGAGAGAAAAGCAACCCUAGAAAGCUUUCCCGUCUAACACACCU**GUAGUCUAGACGGCCAGAGGG5’**

**Supplementary Figure 5. Structure and sequences of exemplary *e*dsRNAs and the control dsRNAs used** (schematically shown in **Figure 4A)**. The sequences of the two strands of the respective dsRNAs are shown. The *e*dsRNAs contain six 21 nt- or 22 nt-long *e*siRNA sequences (guide strands indicated in different colors; see text and **Figure 4A**). The control RNA dsCMV contains a 126 nt-long double-stranded fragment (corresponding to a length of six 21 nt-long siRNAs) derived from CMV RNA 2. The fragment contains, by chance, two overlapping siRNAs that were identified as *e*siRNA candidates in the screening procedure (siR557 and siR540; guide strand sequences underlined and italicized, respectively). The control RNA dsGFP contains a 126 nt-long double-stranded fragment of GFP mRNA including the sequence of the control siRNA siR gf698 used in previous experiments (guide strand sequence in lowercase). All dsRNAs contain pseudo-siRNA sequences at the ends (indicated in bold). The examples shown here generate a 2 nt 3’ overhang at both ends after annealing of the single-stranded components; however, *e*dsRNAs with blunt ends were also generated (see text). The different parts of the *e*dsRNAs dsCMV6-21o or -22o are indicated as follows (guide strand sequences each): Upper strand (5’-3’): bold black, pseudo-siRNA; red, siR1172; yellow, siR1489; brown, siR359. Lower strand (3’-5’): bold black, pseudo-siRNA; dark green, siR380; blue, siR2041; light green, siR1020 (see also **Figure 4A**).

**Supplementary Figure 6**

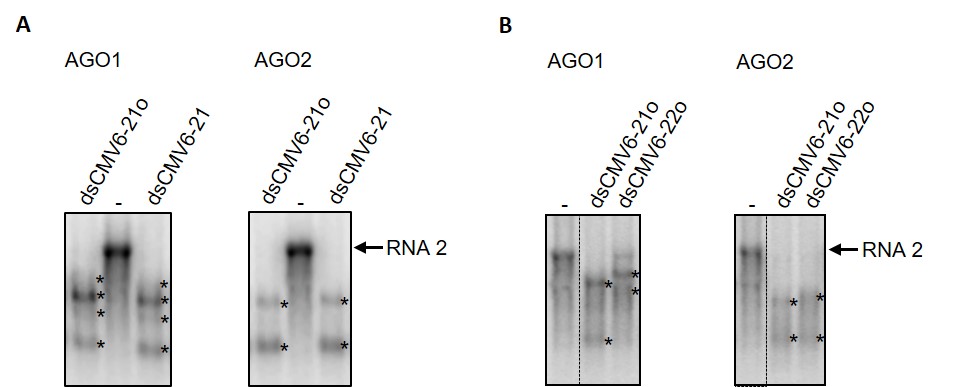

**Supplementary Figure 6. Effect of termini structure and *e*siRNA size on the *e*dsRNA-mediated *in vitro* slicer activity.** AGO1 or AGO2 mRNAs were translated in BYL in the presence of *e*dsRNAs that contained *e*siRNA sequences directed against CMV RNA 2. Thus, AGO1- or AGO2/RISCs were programmed with siRNAs that were processed from the *e*dsRNAs by DCLs present in BYL. ^32^P-labeled, single-stranded CMV RNAs 2 was added as a target, and siRNA-mediated cleavage was analyzed by denaturing agarose gel electrophoresis of total RNA and subsequent autoradiography. **(A)** Results of representative slicer assays performed with *e*dsRNAs having either blunt ends (dsCMV6-21) or 2 nt 3’ overhangs (dsCMV6-21o). The *e*dsRNAs contained six 21 nt *e*siRNAs (three active in experiments with AGO1, three active in experiments with AGO2). (**B**) Results of representative slicer assays performed with *e*dsRNAs containing either six 21 nt *e*siRNAs (dsCMV6-21o) or the six corresponding 22 nt *e*siRNAs (dsCMV6-22o). The latter were obtained by extending the guide strand of the 21 nt *e*siRNAs by one nucleotide. Asterisks (*) indicate the cleavage products.

**Supplementary Figure 7**

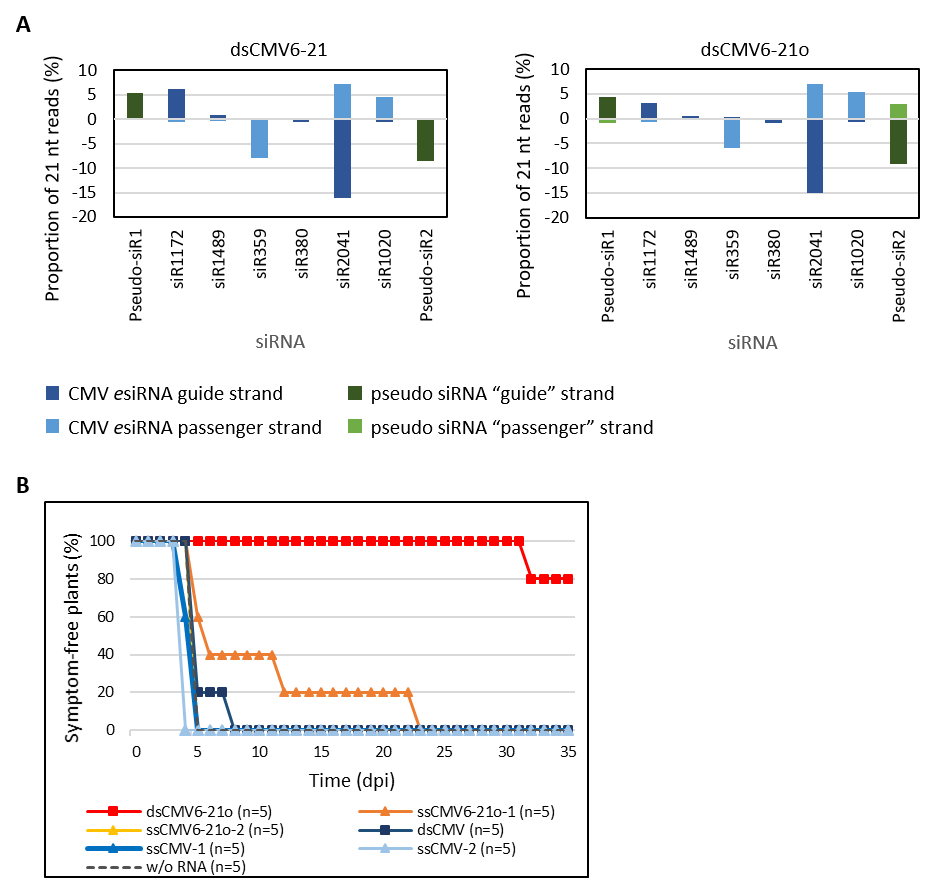

**Supplementary Figure 7. RNA-seq analysis of siRNAs generated in BYL from *e*dsRNAs and comparison of the protective effect with single-stranded RNA. (A)** dsCMV6-21 (blunt ends) and dsCMV6-21o (2 nt 3’-overhangs) were processed in BYL by the endogenous DCLs. Total RNA was isolated, and the small RNA fraction analyzed by RNA-seq (see also **Figure 5**). The image shows the ratio of reads for the guide strand and the corresponding passenger strand for the six CMV RNA 2-targeting *e*siRNAs and the two pseudo-siRNAs on top of each other. In case of dsCMV6-21 there are no mapping 21 nt reads for the “passenger” strands of the pseudo-siRNAs, as the two 3’-nucleotides are missing. (**B**) Comparison of the protective effect of double- and single-stranded RNAs *in planta*. *N. benthamiana* plants were mechanically co-inoculated with the genomic CMV RNAs and the *e*dsRNA dsCMV6-21o, the control RNA dsCMV, or the corresponding single-stranded RNAs (1 and 2) that make up the dsRNAs (see **Figure 4A** and **Supplementary Figure 5**). Inoculation with genomic CMV RNAs served as control. Plants were monitored for the appearance of CMV-specific symptoms for 35 dpi; the image shows the percentage of asymptomatic plants over the entire period of the experiment. Results from one experiment including 5 plants per treatment are shown.
